# Single Cell Sequencing Reveals Glial Specific Responses to Tissue Processing & Enzymatic Dissociation in Mice and Humans

**DOI:** 10.1101/2020.12.03.408542

**Authors:** Samuel E. Marsh, Tushar Kamath, Alec J. Walker, Lasse Dissing-Olesen, Timothy R. Hammond, Adam M.H. Young, Abdulraouf Abdulraouf, Naeem Nadaf, Connor Dufort, Sarah Murphy, Velina Kozareva, Charles Vanderburg, Soyon Hong, Harry Bulstrode, Peter J. Hutchinson, Daniel J. Gaffney, Robin J.M. Franklin, Evan Z. Macosko, Beth Stevens

## Abstract

A key aspect of nearly all single cell experiments is the necessity to dissociate intact tissues into single cell suspensions for processing. While many protocols have been optimized for optimal cell yield, they have often overlooked the effects that dissociation can have on *ex vivo* gene expression changes during this process. Microglia, the brain’s resident macrophages, are a highly dynamic population that are extremely sensitive to their microenvironment and have been shown to dramatically alter their transcriptome upon stimulation. We demonstrate that use of enzymatic dissociation methods on mouse central nervous system (CNS) tissue induces an aberrant gene expression signature in microglia that can significantly confound downstream analysis. To minimize this issue, we developed a flexible protocol, that can be used with existing enzymatic protocols for fresh tissue, to eliminate artifactual gene expression while allowing for increased cell type diversity and yield. We demonstrate efficacy of this protocol in analysis of diverse CNS cell types and sorted myeloid populations while using enzymatic dissociation. Generation of new and reanalysis of previously published human brain single nucleus RNAseq (snRNA-seq) datasets reveal that a similar signature is also present in post-mortem tissue. Through novel snRNA-seq analysis of acutely-resected neurosurgical tissue we demonstrate that this signature can be induced in human tissue due to technical differences in sample processing. These results provide key insight into the potential confounds of enzymatic digestion and provide a solution to allow for enzymatic digestion for scRNA-seq while avoiding *ex vivo* transcriptional artifacts. Analysis of human tissue reveals potential for artifacts in current and future snRNA-seq datasets that will require deeper analysis and careful consideration to separate true biology from artifacts related to post-mortem processes.

## Introduction

The field of single cell and single nucleus RNA sequencing (scRNA-seq, snRNA-seq) has expanded at a rapid rate with the number of studies and numbers of cells profiled increasing dramatically year-over-year^1,2^. However, bio-logical discovery through these technologies is predicated on being able to faithfully measure cell-type-specific expression patterns *ex vivo*. Due to both the effort and cost of performing scRNA-seq, considerable effort has been put toward optimizing protocols to isolate healthy, viable cells from intact tissues. Enzymatic digestion using one or multiple enzymes has become the preferred method for isolating whole cells from many solid tissues due to the ability of proteolytic enzymes to easily digest tough tissue substructures^3–8^. Most of these isolation protocols are performed at elevated temperatures (typically 37°C) to maximize enzyme efficacy and therefore maximize cell yield and viability. However, as the goal of most, if not all, scRNA-seq studies is to produce transcriptional profiles that are reflective of the *in vivo* state it is critical to determine whether an isolation procedure modifies the transcriptional state of cells *ex vivo* before undertaking such studies.

While fresh tissue is easily available from model systems, profiling human tissue often requires the use of frozen tissue. The advancement of snRNA-seq protocols has enabled the profiling of fresh-frozen organs from tissue banks across a variety of conditions^9,10^. Single nucleus sequencing is especially critical for tissues like the brain where obtaining brain tissue from live donors is difficult and only possible under very specific disease or injury conditions. However, few studies have examined the brain for differences in transcriptional states as a result of tissue handling or the post-mortem process^11^ and to our knowledge none have done so in cell-type specific manner using snRNA-seq.

In the current study we sought to understand the cell-type specific transcriptional responses to dissociation of mouse brain tissue with enzymatic digestion at 37°C, and to evaluate how *ex vivo* processing and post-mortem may affect the transcriptional profiles of nuclei isolated from human brain tissue. We find that in mice, microglia are preferentially sensitive to *ex vivo* artifacts, dramatically alter-ing their transcriptome, following enzymatic dissociation. We characterize this *ex vivo* response through an in-depth comparative analysis of multiple microglia isolation protocols. To prevent this aberrant transcriptional state following enzymatic digestion, we optimize a flexible protocol that utilizes the addition of transcriptional and translational inhibitors during multiple steps of the dissociation process. We demonstrate that this protocol effectively eliminates the *ex vivo* activation that accompanies traditional enzymatic dissociation.

Additionally, we perform snRNA-seq on post-mortem human brain, as well as a reanalysis of several published datasets to determine if this aberrant response is observed in post-mortem tissue. We find a similar gene signature is present in post-mortem microglia and astrocytes, across all snRNA-seq datasets analyzed. Through the use of acutely resected neurosurgical tissue, we reveal that a similar signature can be detected in microglia as result of technical variability in sample processing. This result suggests the possibility that the presence of this signature in post-mortem brain samples may be artifactual. Together our results provide a methodological solution for preventing artifactual gene expression changes during enzymatic digestion of brain tissue and a reference for future deeper analysis on the potential confounding states present in post-mortem human samples.

## Results

### scRNA-seq reveals that mouse microglia are highly sensitive to *ex vivo* alterations in gene expression

As the tissue resident macrophages of the brain, microglia are highly sensitive to perturbations in their environment^12,13^. In our previous work, creating a microglia scRNA-seq atlas across the lifespan of mice, we optimized a cold mechanical dissociation protocol to isolate microglia with minimal *ex vivo* alterations^14^. However, the yield using this mechanical protocol is lower than enzymatic-based protocols, especially as mice age, and therefore may not be suitable for all experimental designs.

To systematically characterize the microglial response to enzymatic dissociation we compared our previously optimized cold mechanical dissociation to enzymatic digestion. As we suspected that enzymatic dissociation would induce an aberrant transcriptional signature in microglia, we modified a protocol previously designed to prevent *ex vivo* neuronal activity^15^ and added a cocktail of transcriptional or transcriptional and translational inhibitors during multiple steps of the experimental design **(Figure 1a)**. In addition to the inhibitor cocktails, we maintained tissue/cell solutions on ice at all times with the exception of the enzymatic digestion itself which was performed at 37°C. Following dissociations, microglia were sorted for any dual positive CD45+/ CD11b+ cells **(Figure 1a, SI Figure 1a-b)**. Of note, we observed substantial loss of some cell surface receptors following enzymatic digestion, in line with previous work in peripheral immunology **(SI Figure 1d-f; SI Note 1)**^3–8^. This finding has important implications any study investigating extracellular proteins, but is especially critical for single cell techniques such as flow cytometry or CyTOF, and multi-modal methods (i.e. CITE-Seq)^16,17^ **(SI Note 1)**.

**Figure 1:**
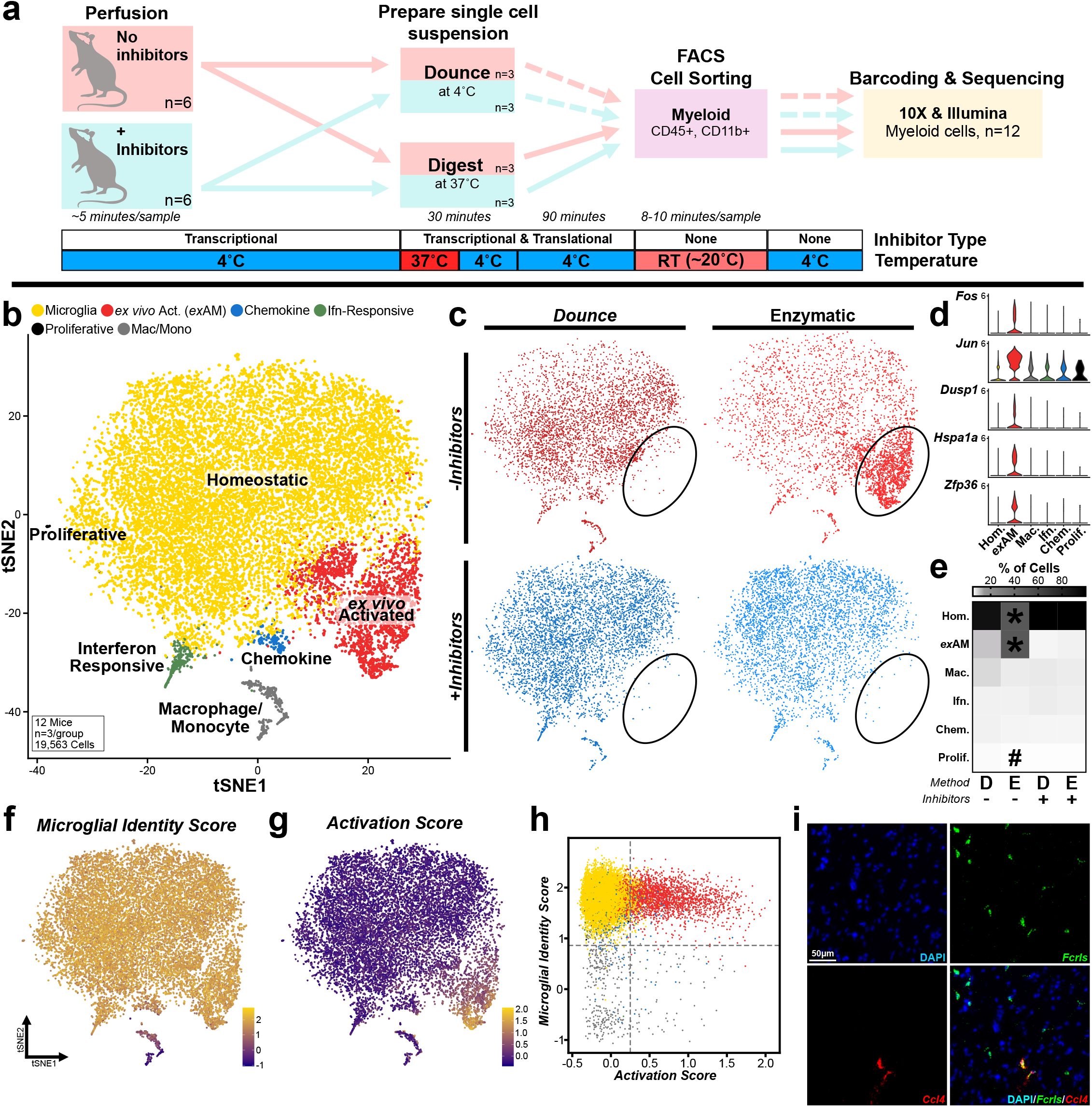
Analysis of sorted microglia confirms profound effect of enzymatic digestion on microglial gene expression via single cell sequencing. **a.** Experimental design schematic for sorted mouse myeloid cells scRNA-seq experiment **(see Methods and SI Figure 1a-c). b**. tSNE plot for the 19,563 cells from n=12 mice (n=3/group) colored and annotated by cluster **(see SI Figure 2). c**. tSNE plot split by experimental subgroup highlights enrichment of *ex*AM cluster in ENZ-NONE group. **d**. Heatmap of the mean percent of cells in each cluster across conditions (**SI Table 3)** (* FDR<0.005 for ENZ-NONE vs. all other groups; ^#^ FDR < 0.05 DNC-NONE vs. ENZ-INHIB; Benjamini & Hochberg correction for multiple comparisons, **See Methods: Differential Abundance Testing**). **e**. Gene expression of several *ex*AM cluster markers across each cluster. **f-g**. Visualization of gene module scoring results plotted on tSNE coordinates. **f**. Microglial identity score **(See Methods & SI Table 4). g**. Activation score based on consensus DE genes from ‘Meta-Cell’ pseudobulk analysis **(See Methods & SI Tables 4-5). h**. Plot of microglial identity score vs. activation score colored by cluster annotation from panel **b. i**. smFISH using RNAscope for microglial marker *Fcrls* (green), cluster marker *Ccl4* and counterstained with DAPI. Scale bar = 50 μm.

Following QC, we analyzed 19,563 cells from 12 mice (n=3 mice per condition). In addition to a small number of brain-/ brain-border-associated macrophages/monocytes (Mac/Mono), we identified 4 primary clusters of microglia that were present in all four groups **(Figure 1b)**. In line with our previous work^14^, we find that adult microglia are largely homogenous with the majority of cells (∼81%) falling into what we term the “homeostatic” cluster. We also identified several smaller clusters of microglia, which we annotated on the basis the cluster-specific marker genes: proliferative (i.e. *Top2a, Birc5*), interferon-responsive microglia (i.e. *Ifitm3, Irf7*), and chemokine-expressing (i.e. *Ccl3, Ccl4*) **(Figure 1b, SI Figure 2)**.

We also identified an additional cluster, nearly exclusively composed of cells from samples digested with enzymes and no inhibitors (ENZ-NONE) **(Figure 1c-d)**, which was characterized by expression of several immediate early genes (IEGs) and stress-induced genes (i.e. *Fos, Jun, Hspa1a, Dusp1*) **(Figure 1e, SI Figure 2c-d)**. Given the nature of the genes expressed and that its overwhelming enrichment for the ENZ-NONE group, we termed this cluster *ex vivo* “Activated” Microglia (*ex*AM). Statistical analysis of contribution of each group to each cluster, found that the ENZ-NONE samples exhibited significant decreases in the proportion of cells in the homeostatic cluster and increased proportion of cells in the *ex*AM cluster compared to all of the other groups (*FDR 0.004; ENZ-NONE vs. all other groups)**(Figure 1d, SI Table 3)**. While the proliferative cluster was statistically significant between the DNC-NONE and ENZ-INHIB groups, that result is difficult to interpret as the proliferative cluster was only 0.1% of total cells (27/19,563) across all samples.

To identify the differentially expressed genes (DEGs) following enzymatic digestion we performed sample-level comparisons^18^ by creating pseudo-bulk “Meta-Cells”^14^ by aggregating expression across all cells within each bio-logical replicate and then performed differential expression (DE) analysis using DESeq2^18,19^. We identified a consensus DE signature of upregulated genes that was shared among the comparisons of each experimental group to the ENZ-NONE group (**SI Table 4-8)**. This signature was composed of a number of different types of genes including IEGs (i.e. *Fos, Jun*), genes induced by cellular response to stress (i.e. *Hspa1a, Dusp1*) ^20–22^, genes associated with regulation of transcription (*Hist1h1d, Hist1h2ac*), chemokine genes (*Ccl3, Ccl4*), and parts of the NF-κB signaling cascade (*Nfkbiz, Nfkbid*) **(SI Table 4-5)**. To visualize which cells in the dataset were enriched for this signature, we performed gene module scoring^23^. We created two scores, one using a core micro-glial gene signature **(Figure 1f, SI Table 4)** aggregated from previous publications and another score, which we refer to as the “activation” score, using the consensus DEG list **(Figure 1g-h)**. Analysis of enrichment of this “activation” signature plotted both on tSNE coordinates **(Figure 1g)** and against the microglial identity score **(Figure 1h)** demonstrated that this activation signature was almost exclusively enriched in cells from the *ex*AM cluster.

The overlap of two cluster markers (*Ccl3, Ccl4*) between the chemokine and *ex*AM clusters required additional examination to confirm that the small chemokine population was truly an *in vivo* state. We performed single molecule *in situ* hybridization (smFISH) on acutely isolated and snap-frozen tissue, which preserves cells in as close to *in vivo* state as possible. In line with our previous study and others ^14,24^, we confirm that, while extremely rare, the chemokine cluster *Ccl4+* cells are a true *in vivo* state and not merely an additional dissociation-induced response **(Figure 1i)**.

To examine whether presence of inhibitors had any negative consequences we performed further DE comparisons. Analysis of DNC-NONE vs. DNC-INHIB groups found no significant DEGs between the groups **(SI Table 9, 12)**, indicating no negative effect of inhibitors. Further analysis between the other groups found that there may be a small subset of genes that are modulated by the presence of inhibitors **(SI Figure 3, SI Table 8-12)**. However, the magnitude of the changes were very small and only a handful of genes were affected. These results indicate that overall, the presence of inhibitors appears to have negligible adverse effects on gene expression in the current study.

### Presence of artifactual microglia gene expression signature is not specific to enzyme type, dissociation protocol, or sequencing technology

To confirm that the *ex*AM signature was not specific to our protocol or the scRNA-seq technology used in the present study, we re-analyzed previously published microglial/CNS datasets for the presence of the *ex*AM signature^25–29^. This re-analysis included datasets processed with several different scRNA-seq technologies (10X Genomics 3’ v1, 10X 3’ v2, Smart-Seq2, Drop-Seq, and Microwell Seq), dissociation enzymes (papain, collagenase), brain regions (whole brain, cortex, SVZ, spinal cord), and various other differences in exact digestion method, timing, or other parameters **(SI Figure 4a-g)**. In all 5 of these datasets, we observed significant enrichment of *ex*AM signature via gene module score enrichment in the myeloid populations consistent with *ex vivo* activation **(SI Figure 4a-e)**. We also reanalyzed our previous study^14^, that utilized cold Dounce homogenization and found minimal presence of cells enriched for the *ex*AM signature **(SI Figure 4f)**, confirming proper cold Dounce homogenization is sufficient to prevent *ex vivo* activation. Finally, analysis of data from a separate mouse experiment in our lab revealed the importance of the inclusion of true biological replicates in scRNA-seq experiments, as artifacts are possible even when using ideal isolation methods when there are small differences in sample processing prior to cell capture **(SI Figure 5; SI Note 2)**.

### Artifactual gene expression signature is shared by other immune populations and tissues

We also wondered whether other myeloid lineage CNS cells were sensitive to this aberrant *ex vivo* activation. Despite the relatively small number of CNS-associated macrophages present in our dataset we were still able to identify an enrichment of the artifactual *ex*AM signature in these other myeloid cells, again only from animals that were enzymatically digested without inhibitors **(SI Figure 6a-d)**.

These results are in agreement with a previous report in the literature which found that dural and choroid plexus macrophages also appear to exhibit a dissociation-induced signature^30^. This study identified a list of greater than 200 genes associated with dissociation in their dataset. Our consensus *ex*AM signature does overlap with the dissociation signature in Van Hove et al., (19 of 25 genes in consensus *ex*AM list **(SI Table 4-5)** were also present in the Van Hove List; **SI Table 13a**), but many of the genes they identified do not exhibit shifts in our dataset. We also examined two other studies in the literature which have utilized scRNA-seq to examine dissociation induced genes in non-CNS tissue (mouse muscle and kidney)^31,32^. Comparison of our consensus *ex*AM gene list to those studies yielded overlap of 16 and 11 genes, respectively **(SI Table 13b-c)**. When we compare the three lists of overlapping genes with our study, we find the majority of genes are shared across all tissue/cell types indicative of a common cellular response to dissociation **(SI Table 13d)**. However, we do find some genes that appear tissue/cell-type specific to myeloid cells, such as *Ccl3* and *Ccl4* **(SI Table 13d)**.

### scRNA-seq reveals that mouse microglia are especially sensitive to *ex vivo* alterations in gene expression compared to all other CNS cell types

While our cold Dounce mechanical protocol works well for microglial isolation, all other CNS cell types are lost. In order to analyze all CNS cell types simultaneously via scRNA-seq, while maintaining good cell viability, enzymatic dissociation is required. However, few studies have examined whether CNS cell types other than microglia/myeloid cells exhibit altered gene expression when isolated from their *in situ* environment^33–35^.

To characterize the transcriptomic response of all CNS cell types following enzymatic digestion, we used the same transcriptional/translational inhibitor cocktail that we used for our analysis of microglia with small changes (**Figure 2a)**. We analyzed 10,166 cells (post-QC; n=2/group) and identified 16 broad clusters, representing all of the major cell types found in the CNS and a very small cluster of contaminating red blood cells **(Figure 2b, SI Figure 7a, 7c)**.

**Figure 2:**
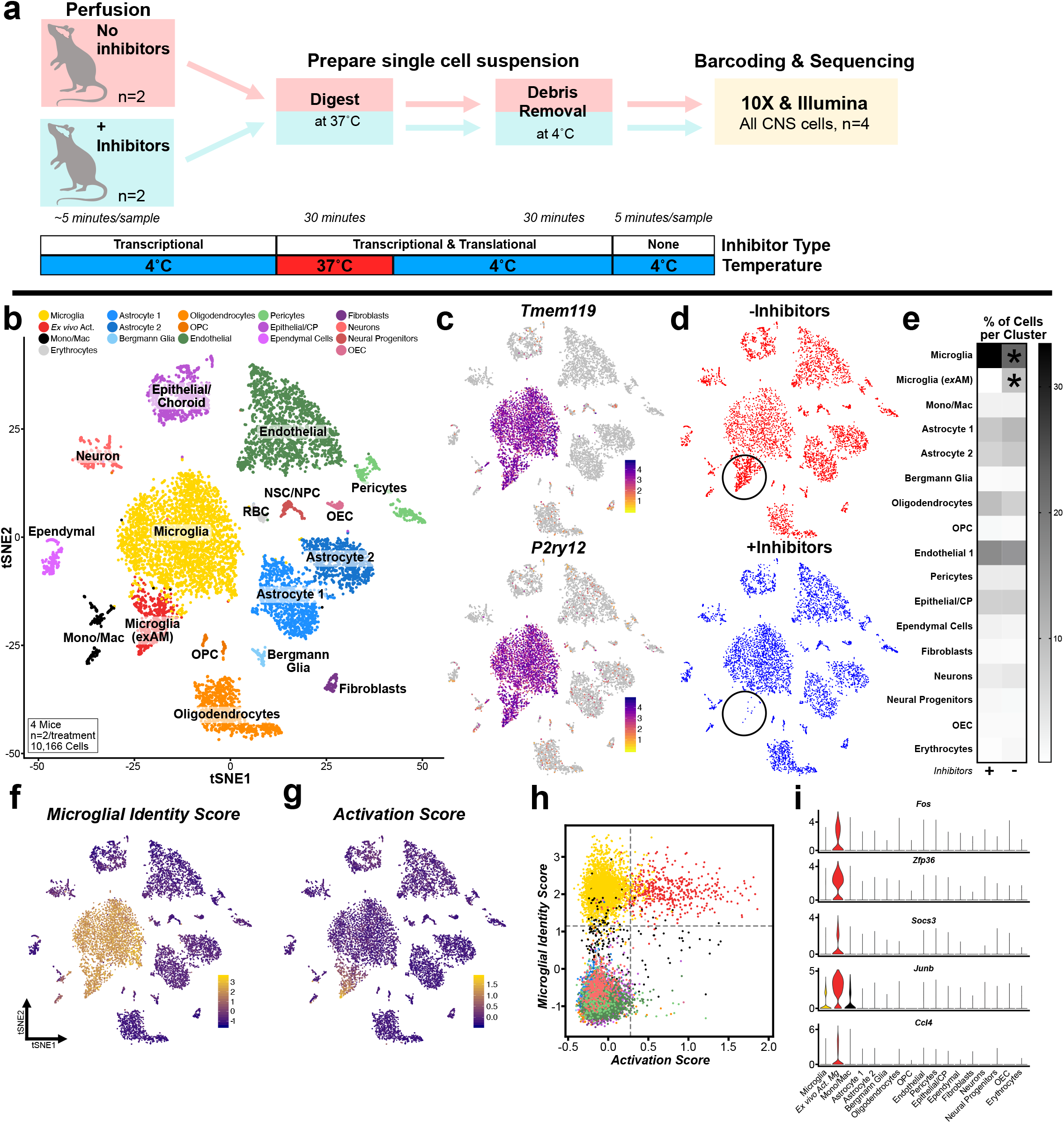
Enzymatic dissociation induces cell-type specific artifactual gene expression in mice. **a.** Experimental design schematic for scRNA-seq of all CNS cell types **(see Methods). b**. tSNE plot of 10,166 cells from 4 mice (n=2/group), annotated and colored by cell type **(see SI Figure 7). c**. Expression of microglial specific genes *Tmem119* and *P2ry12* clearly defines two microglial clusters. **d**. tSNE plot split by experimental group (+/-inhibitor cocktail) highlights lack of *ex*AM cluster in samples digested with inhibitors present (bottom; circled). **e**. Heatmap displaying mean percent of cells in each cluster across conditions (* FDR<0.0005; Benjamini & Hochberg correction for multiple comparisons, **See Methods: Differential Abundance Testing**). **f-g**. Gene module scoring results plotted on tSNE coordinates. **f**. Microglial identity score **(See Methods & SI Table 4). g**. Activation score based on DE genes from ‘Meta-Cell’ pseudobulk analysis **(See Methods & SI Table 4, 15). h**. Scatterplot of gene module scores from **f** and **g** colored by cluster from panel **b. i**. Enrichment of genes from activation score in *ex*AM microglia cluster.

Microglia were the most prominent cell type in the dataset using this digestion protocol **(Figure 2b-c, e)**, likely indicating that the Miltenyi dissociation system utilized is not appropriate for general unbiased profiling of brain tissue when more accurate cell proportions are desired. Plotting the cells by group, revealed one cluster of microglia that was almost completely composed of cells digested without inhibitors present **(Figure 2d, circled)** and expressed multiple *ex*AM markers **(SI Figure 7c)**. Analysis of the proportion of cells from each group in each cluster revealed that only the microglia clusters differed significantly between the groups (*FDR < 0.003) **(Figure 2e; SI Table 14)**. To determine the DEGs between the groups, we again performed sample-level comparisons using our pseudobulk DESeq2 pipeline. This DE analysis revealed a signature of 18 genes that were significantly upregulated (padj < 0.05, log_2_fc > 0.58) and no downregulated genes when cells were isolated without inhibitors **(SI Table 4, 15; Figure 2i; SI Figure 7c)**. These DEGs exhibited significant overlap with the previously identified DEG list from the microglial analysis confirming that enzymatic digestion and not FACS was the key factor in the induction of this aberrant response. We again performed module scoring to visualize the enrichment of this signature. Analysis of enrichment of the *ex*AM signature demonstrated that the signature was present almost entirely in cells from the *ex*AM cluster **(Figure 2f-h)**. The enrichment of this signature can also be seen in the near exclusive expression of many of the DE genes in the *ex*AM cluster **(Figure 2i)**.

To perform a deeper analysis by cell class and determine if any other cell types exhibited significant effects following dissociation, we performed subclustering analysis. Subclustering revealed that microglia/myeloid cells were the only major cell class that exhibited clustering driven by the presence/absence of the inhibitor cocktail **(SI Figure 8a-f)**. However, in other cell types we did identify increased IEG expression in cells digested without inhibitors. For instance, in analysis of oligodendrocyte/ oligodendrocyte precursor cells, we did observe slight increases in either expression level or number of expressing cells for some of the DEGs, when inhibitors were absent **(SI Figure 8g-i)**. These results are in agreement with previous *in vitro* work which found that microglial reaction to heat stress was dramatically faster than other CNS cell types^36^. This result indicates that *ex vivo* artifacts occurring in other CNS cell types is possible and that caution should be taken in a dataset-specific manner as differences in digestion protocol could lead to activation in non-microglia/ myeloid CNS cell types.

### Analysis of human post-mortem snRNA-seq identifies similar gene signatures in both microglia and astrocytes

Following our characterization of mouse tissue, we hypothesized that post-mortem processes and/or post-mortem interval (PMI) might also induce a similar artificial signature in human tissue. We performed nuclei isolation, sorting, and snRNA-seq using the 10X 3’ V3 gene expression kit, from 3 post-mortem donors with a wide spread of PMIs (**Figure 3a)**. Integrative analysis using LIGER^37,38^, of 47,505 nuclei across all 3 donors, identified all of the expected major cell types present in the CNS, as well as some contaminating peripheral immune cells **(Figure 3b-c; SI Figure 9a-c)**.

**Figure 3:**
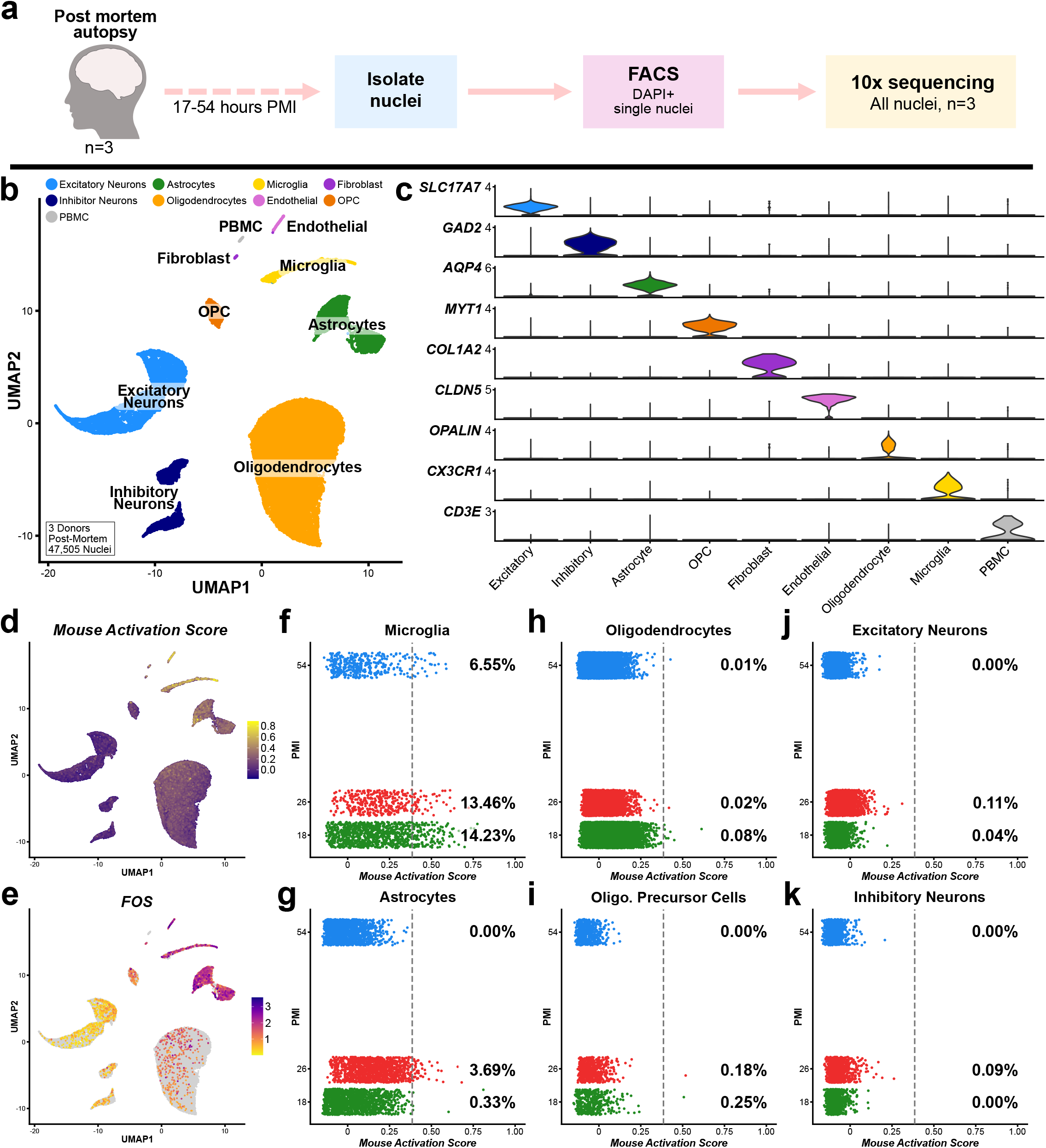
snRNAseq of human post-mortem tissue identifies enrichment of mouse dissociation gene signatures in microglia and astrocytes. **a.** Experimental design schematic for snRNAseq of all cell types from frozen post-mortem brain tissue. **b**. UMAP plot of 47,505 nuclei via snRNAseq from 3 post-mortem subjects following LIGER analysis, colored by major cell type. **c**. Expression of canonical marker genes delineates major cell types. **d**. Visualization of the gene module scoring of mouse DEG signature on human post-mortem snRNAseq dataset. **e**. Gene expression of activation signature gene *FOS* across clusters. **f-k**. Plot of mouse activation score vs. sample PMI for each of the major CNS cell classes present in the dataset. Percentages denote number of nuclei above enrichment threshold denoted with gray dotted line.

To determine whether the *ex vivo* dissociation signature we identified in mice was enriched in any particular cell type in human post-mortem data, we performed gene module scoring (using the DEG set identified in the analysis of all CNS cell types in the mouse) **(SI Table 4, 15**). Similar to mice, human post-mortem microglia exhibited significant enrichment of this signature, as did a small number of astrocytes **(Figure 3d-e)**.To quantify the enrichment of the signature across the various cell types we plotted the percent of nuclei above the enrichment threshold (**See Methods)** vs. PMI for each of the donors **(Figure 3f-k)**. This analysis demonstrated that microglia and some astrocytes exhibited the greatest enrichment of the mouse signature. We also found that the percent of nuclei above threshold per sample was both highly variable and did not correlate with PMI (**Figure 3f-k)**.

To increase the power of our analysis examining this potentially confounding signature, we performed reanalysis of several published snRNA-seq datasets (**SI Table 2**)^39–41^ using our LIGER-based pipeline. Across all of the datasets we analyzed 49 samples and 248,026 nuclei and included both “control” patients and donors with Alzheimer’s disease **(SI Figure 9)**.

When we performed the same module scoring using the mouse signature, we found that microglia and some astrocytes were again the most enriched for the signature **(SI Figure 9 rightmost column)**. Due to differences in gene/transcript detection sensitivity across these datasets, comparison of the raw expression values is not appropriate **(SI Figure 9; SI Note 3)**. While these results indicated some enrichment in post-mortem tissue, we also wanted to know if we could identify this signature in post-mortem data without *a priori* knowledge of what genes were a part of the signature. One advantage of LIGER’s NMF-based analysis is that it generates a series of “factors” that are shared across the datasets present in the analysis. These shared factors often identify gene signatures that correspond to biologically relevant signals^37^, such as genes that define particular cell type/ state or genes that are part of a particular biological pathway. To determine whether we could identify a similar signature in the human data, we first subsetted each of the major cell types from each dataset and then performed a combined subclustering analysis with LIGER. In total we found a factor in each cell type that shared at least 1 gene with the combined mouse signature gene lists **(SI Table 16**). We examined the degree of overlap between the thresholded genes in each of these shared factors (**See Methods**) and the combined mouse gene lists. Only the factors from the microglial and astrocyte subclusterings exhibited a 25% or greater overlap with the mouse gene list **(SI Table 16)**.

The microglia post-mortem factor (**Figure 4a)** was strikingly similar to the mouse signature with 79% (30/38) of the genes directly overlapping with mouse signature or were genes from similar families/pathways. The astrocyte post-mortem factor **(Figure 4b)** exhibited modest overlap with the mouse signature as 44% (12/27) directly overlapped or were genes in similar families. Finally, there were 3 genes that overlapped between the microglia and astrocyte factors that were not a part of consensus mouse gene lists (*UBC, DDIT4*, and *HSPB1*), but which unsurprisingly are also part of cellular stress and damage response machinery^42–45^.

**Figure 4:**
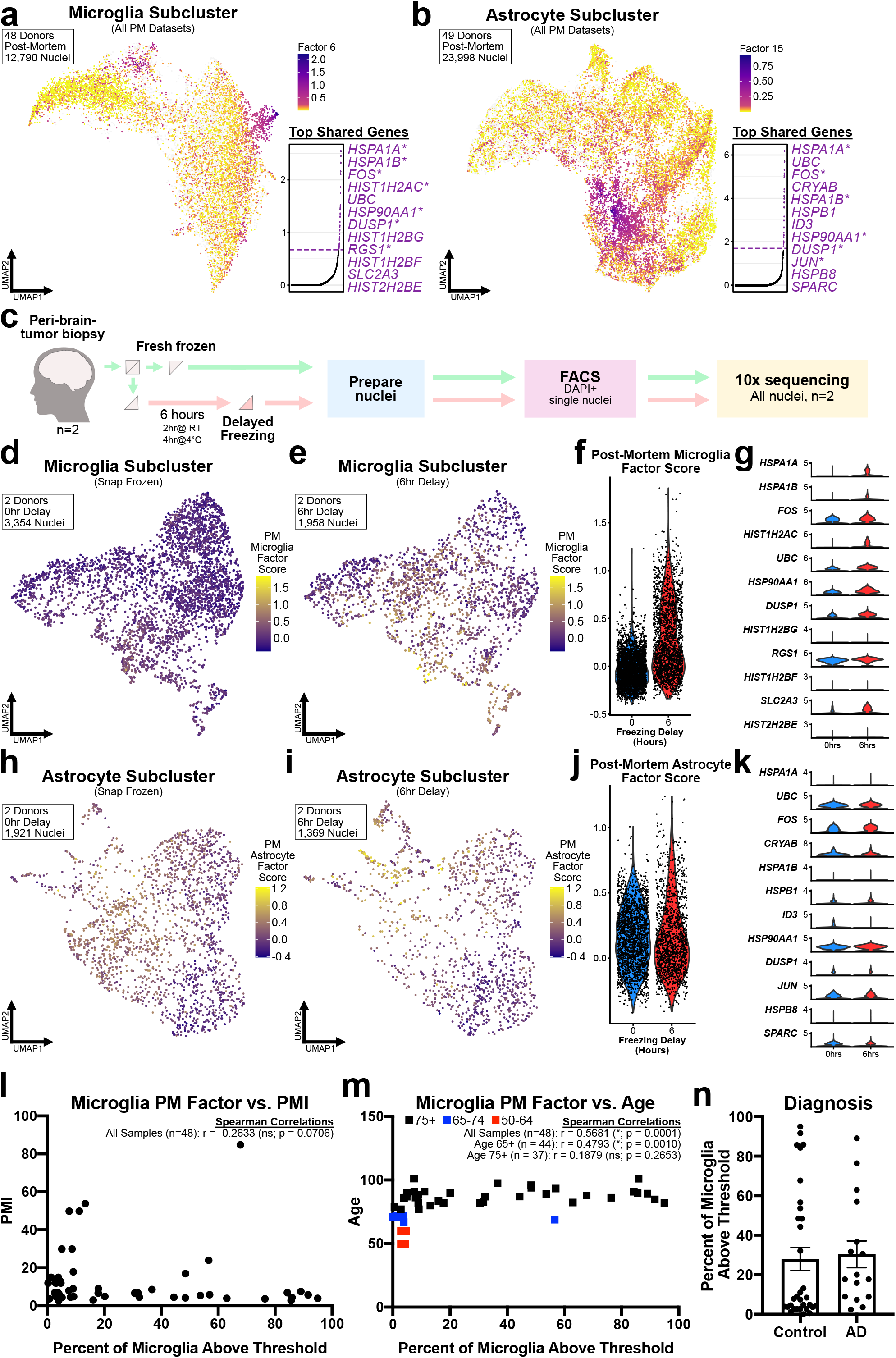
LIGER analysis independently identifies similar gene expression signatures in post-mortem data that are enriched in microglia following altered sample processing. **a.** UMAP plot visualizing the enrichment of shared LIGER factor for 12,790 microglial nuclei from 48 samples across all datasets. **b**. UMAP plot visualizing the enrichment of shared LIGER factor for 23,998 astrocyte nuclei from 49 samples across all datasets. For both **a-b** inset displays plot of normalized cell-specific factor loading scores across all genes in dataset (Dashed line indicates threshold cutoff for top genes for downstream analysis; **SI Tables 16**). Top loading genes in order are shown to the right of inset plot. **c**. Experimental design schematic for experiment to analyze the effects of altered sample processing on gene expression. **d-f**. Visualization of gene module scoring results for score based on post-mortem microglia factor, from **a**., in both snap frozen and 6 hour delayed freezing microglia nuclei **d-e** on UMAP coordinates or **f** via violin plot split by experimental group. **g**. Gene expression of top 12 loading genes in microglial factor from **a** split by experimental group **(See SI Table 17 for DEG results). h-j**. Visualization of gene module scoring results for score based on post-mortem astrocyte factor, from **b**., in both snap frozen and 6 hour delayed freezing astrocyte nuclei **h-i** on UMAP coordinates or **j** via violin plot split by experimental group. **k**. Gene expression of top 12 loading genes in astrocyte factor from **b** split by experimental group **(See SI Table 18 for DEG results). l-m**. Correlation of PMI and age of donor vs percent of microglia above score threshold for microglia factor score in each post-mortem sample, graph annotations list Spearman r values and significance. **n**. Plot of percent of microglia above score threshold for microglia factor score in each post-mortem sample split by diagnosis.

### Technical variation in sample processing of acutely-isolated human tissue is sufficient to induce to induce similar microglial but not astrocyte gene signatures

In order to examine whether or not the presence of this signature in human microglia and astrocytes may represent a confounding technical aspect rather than a true *in vivo* state, we performed snRNA-seq analysis of acutely-isolated human brain tissue. Peri-tumor tissue, normally discarded during surgery, was obtained and divided into 2 fractions. Half was immediately snap-frozen in liquid nitrogen and the other half was placed in an artificial cerebrospinal fluid (aCSF) medium and incubated for 2 hours at room temperature and another 4 hours at 4°C before being snap-frozen **(Figure 4c)**. This protocol was intended to test whether altered tissue processing, unrelated to digestion, could induce this artificial signature.

We then performed LIGER-based integrative analysis and subsetted the microglia and astrocytes for further analysis. To determine whether a technical variable (freezing delay) induced a similar signature to that observed in post-mortem nuclei, we performed gene module scoring using the LIGER factor gene lists from post-mortem microglia or astrocytes as input **(SI Table 16)**. Analysis of microglia nuclei found significant enrichment of the putative “artificial” microglia factor at the 6hr timepoint compared to 0hr timepoint **(Figure 4d-f)**. Analysis of the individual genes in the signature also found that 31/38 were statistically significantly enriched at the 6hr timepoint (**Figure 4g; SI Table 17)**. These results suggest that the microglial LIGER factor we identified can be induced by *ex vivo* gene expression changes induced as result of a number of cell-autonomous molecular processes of likely including, among others, hypoxia, cell death, and temperature stress.

Similar analysis of the astrocytes using the post-mortem astrocyte factor revealed no enrichment pattern at the 6hr timepoint (**Figure 4h-j**). Analysis DEGs also found that only 4/27 genes from the LIGER factor were differentially expressed and of those only *ATF3* overlapped with the mouse signature **(Figure 4k; SI Table 18)**. However, as previously discussed with the mouse results, the observation of a response in microglia but not in astrocytes may simply be due to the different speeds at which different CNS cell types respond to stress^36^.

### Human post-mortem microglial gene signature does not correlate with known metadata variables

Given that our results, that the post-mortem microglia LIGER signature could be artificially induced, next we wanted to determine whether enrichment of that signature correlated with PMI or other metadata variables. To perform this analysis, we quantified the proportion of microglia or astrocyte nuclei with a gene module score above the enrichment threshold (**See Methods)** per sample in each of the datasets. Comparing to metadata variables across the datasets the only statistically significant correlation was found with age of the donor **(Figure 4l-n)** (Spearman: r = 0.3208, *p = 0.0001, n = 48). However, this correlation appears to be strongly driven by a small number of samples from younger donors from a single study ^40^ **(Figure 4m; red and blue squares)**. Analysis of correlation only in samples ages 75+ found no significant correlation (Age 75+ Spearman: r = 0.1879, *p = 0.2653, n = 37). Similar analyses across the postmortem astrocytes found nearly identical results to microglia analysis **(SI Figure 11a-c)**. Future analyses of a greater number of middle-age samples will be needed to confirm whether these younger samples are simply anomalous or whether the age correlation across the entire dataset can be replicated.

## Discussion

The ability to profile entire organs and tissues at a single cell resolution has the potential to transform our under-standing of both normal and disease biology^46–49^. Further advances in multimodal profiling, linking various combinations of transcriptomics, proteomics, epigenomics, and spatial location will only serve to enhance the insights that will be gained from this invaluable technology^50,51^. However, these biological discoveries rest on being able to faithfully measure cell-type-specific expression patterns *ex vivo*. It is critical that best practice protocols be established in this emerging field, so that reliable data, that as closely reflects the *in vivo* cellular state as possible, is obtained. We therefore undertook the present study to both characterize issues that occur during tissue dissociation or post-mortem processes in the mouse and human brain and find novel solutions to aid future studies.

While our previously optimized cold Dounce homogenization works very well for microglia under most conditions, it does have its limitations. While we were still able to recover sufficient numbers of cells for scRNA-seq even in very old mice^14^, the need for greater yield, especially for older or less sensitive approaches, is likely why many widely-used protocols for microglial isolation included enzymatic digestion^52–54^. The other major draw-back to mechanical dissociation is that while it purifies microglia, it leads to loss of other CNS cell types. Consequently, enzymatic dissociation is required in order to take a more holistic approach and study all CNS populations at the single cell resolution. However, emerging results from other tissues/cell types, have begun to characterize dissociation artifacts following traditional enzymatic digestion methods^30–32,55^. One previous study had examined the brain; but the lack of biological replicates and low numbers of cells belonging to rarer cell types, such as microglia, made a detailed characterization of the dissociation signature difficult^34^.

Consistent with the reports from other tissues in mice, we found that typical enzymatic digestion of brain tissue induced host of significant stress-, inflammatory-, and IEG-related responses ^30,31,55^, although interesting we found this induction preferentially in microglia and other brain myeloid cells. We also confirm through reanalysis of the CNS literature, that this aberrant *ex vivo* response is not specific to the enzyme, sequencing technology, or specifics of the protocol utilized in the current study. Under the conditions of the present study the induction of an *ex vivo* induced signature only occurred in microglia and other brain-associated myeloid cells and while not significant, we did find hints that other cell types may show initial signs of this stress/activation response, which may become significant under different conditions. This lessened and/or slower response of other cell types is in line with previous work demonstrating that the degree and speed of cellular response varies across different CNS cell types, with the microglial response being among the largest and fastest^36^. Therefore, it is possible that under different conditions of other enzymatic dissociation protocols (i.e. length of time, enzymes used, additional room temperature steps, etc) that other cell types may also exhibit this signature more prominently.

Through the use of transcriptional and translational inhibitors added at multiple steps of the protocol we were successfully able to eliminate the *ex vivo* activation that occurred in enzymatically digested samples, both in FACS-sorted myeloid cells or a preparation of all CNS cell types. We also find little to no aberrant responses induced by the presence of these inhibitors. We believe that our protocol using this inhibitor cocktail offers much greater experimental and tissue/cell type flexibility than previously developed methods for minimizing the effect of this signature on resulting data. Some recent studies have developed entirely new digestion protocols using cold-active enzymes, such as those from *B. Licheniformis*, which has been successfully applied to the dissociation of mouse kidney^32,55^. However, modifying already functioning protocols, that have often been optimized over years of use, to incorporate a different enzyme cocktail that is cold-active is not necessarily a simple swap. It requires lots of optimization which is often, difficult, time-consuming, and it may never work as well as enzymes that function optimally at near 37°C. Additionally, our attempts to adapt these protocols to brain tissue have been hindered by lack of cold-active DNase^56–58^, which is key component of enzymatic digestion cocktails. By contrast, our inhibitor protocol should be easily adaptable to a variety of dissociation protocols without requiring time consuming testing of new enzymes, simply requiring the addition of the inhibitor cocktail at several steps of the dissociation process regardless of the enzyme being used.

We also believe that this experimental method is superior to proposed computational solutions to this issue. For example, previous studies have sometimes used a computational approach to exclude or regress genes from the clustering steps in the analysis^59^. However, use of this method for dealing with the dissociation signature is based on assumption that the tissues/cells being profiled are not expressing this signature as a result of the inherent biology of a particular condition, age, treatment, genotype, or other important variable. Two recent studies, using a combination of scRNA-seq, spatial transcriptomics, and immunohistochemistry demonstrated that a signature very similar to the dissociation/ activation signature we describe is expressed *in vivo* across a wide range of cancer tissues and is not the result of tissue dissociation^23,60^. Therefore, we believe that removal of these genes from clustering and/or regression of this gene signature should not be performed, as it adds unnecessary bias to the analysis which may cause real biological signals to be unintentionally regressed and go undetected.

Finally, for mouse experiments, while snap-freezing tissue and isolating nuclei on ice does avoid this *ex vivo* activation, it does come with caveats. Specifically, cell-type specific enrichment in unfixed nuclei has proven difficult, leading to the need to sequence very large numbers of nuclei in order to get sufficient numbers of microglia and other rare cell types for properly powered analyses. By contrast, our protocol allows for the use of either unbiased profiling or the use of enrichment/depletion strategies via FACS to increase proportions of rare cell types, though there are caveats to these methods as well (**SI Note 1)**.

Our mouse results have several important implications for the interpretation of previous microglial literature. First, is the need for a reevaluation of studies which have highlighted genes that are a part of our *ex*AM signature as being of particular importance in microglia. For instance, among the genes nearly exclusively expressed following enzymatic digestion without inhibitors were transcription factors such as *Fos, Jun*, and *Egr1*. Previous work, also using enzymatic digestion and room temperature centrifugation, implied that these transcription factors were an important aspect of the microglial post-natal maturation program and were highly expressed in homeostatic adult microglia^61–64^ and a recently published study indicated that lack of induction of this gene signature in MHC II knockout mice was reflective of microglia remaining in an immature, fetal-like, state^65^. Studies using the Ribo-Tag/TRAP-seq based approaches did dispute these results, however, they subsequently implied that induction of these genes was the result of the use of FACS to sort the cells as final step in the isolation, and not the digestion itself^66,67^. However, as we demonstrate in the current study, either cold mechanical dissociation or enzymatic digestion with inhibitors followed by FACS does not induce this signature and the signature was present in our analysis of all CNS cell types which did not undergo FACS, indicating that it was most likely the enzymatic digestion that was used in the analyses from the TRAP-seq studies that likely induced the *ex*AM signature^66,67^. Our findings are also in line with another recent study, which used smFISH to demonstrate that *Fos* and *Egr1*, while expressed in cells adjacent to microglia, were not expressed in microglia at rest^24^.

Finally, recent work has begun to provide new insights into understanding disease-associated microglia signatures^64,68^ that had been previously identified with bulk sequencing methods^69,70^. However, without a proper baseline, the identification of these disease signatures may be impacted and lead to issues in evaluating differentially expressed genes that are then attributed to *in vivo* biology^64^.

The advent of snRNA-seq has opened a wealth of new avenues in neuroscience research through the ability to profile fresh frozen human brain^9,10^. However, there are important quality control assessments which need to be performed and considered, especially as it regards the impact of post-mortem processes^11^. Analysis of our own post-mortem snRNA-seq dataset, as well as reanalysis of literature datasets, revealed a signature similar to the mouse *ex*AM signature, exists in both microglia and astrocytes from human post-mortem studies. This signature is highly variable across donors and was not correlated with PMI. To determine whether this signature could be upregulated in human cells, we designed a study using an artificial technical variable (freezing delay) in acutely-resected neurosurgical tissue. We found that microglia, but not astrocytes, upregulated the post-mortem signatures in the timeframe examined. It seems most likely that this lack of induction in astrocytes in our artificial experiment is related to differences in response time between microglia and astrocytes to physiological stressors^36^. We also demonstrate that while microglia and astrocyte post-mortem signatures correlates with age across the entire dataset, the correlation was no longer significant when restricting the analysis to those age 75+. Given the current number of younger samples in a single dataset, and their origin from a single study^40^, we cannot say for certain whether the overall correlation with age is real or whether the donors below the age of 75 were simply aberrant samples. However, the between subject variability of this signature across samples age 75+ clearly demonstrates that more samples from younger ages are necessary before drawing any conclusions about this correlation.

Importantly in our combined original and literature reanalysis we included samples from both “control” patients and those diagnosed with Alzheimer’s disease (n=32/17; Ctrl/AD). Given the results in other disease states, such as cancer, where part of this activation signature is actually reflective of disease^23,60^, we wanted to examine whether or not this signature was potentially associated with AD. Our analysis revealed no significant differences in these post-mortem signatures between AD and control patients for either astrocytes or microglia. The high between-sample heterogeneity that we and others observe with data from human subjects, highlights the need for scRNA-seq/snRNA-seq analyses in human patients with much greater numbers of samples than most datasets currently contain.

For now, the results of our analysis of human snRNA-seq leads us to caution overinterpretation of biological significance when observing this signature in post-mortem human data. As a greater number of high-quality human datasets become available, this systematic approach and methods will enable better characterization of this signature and the factors that contribute to it. Most currently available human snRNA-seq datasets are limited by both the proportion of microglia per sample and dramatically lower sensitivity in earlier single cell chemistries^71^ **(SI Note 3)**. New datasets with optimized protocols, greater numbers of nuclei per patient, and greater numbers of patients will be critical to resolving both this potentially aberrant state, as well as the *in vivo*, biologically relevant cell states of the human brain^72^. Finally, it is also critical for new studies to provide as much metadata as possible when publishing datasets. Greater amounts of metadata will be critical in our interpretation and understanding of the complexities of human biology with scRNA-seq-/snRNA-seq-based methods.

In conclusion, we provide a single cell method by which to eliminate the aberrant activation when enzymatic digestion is required, that is easy to implement and should require minimal effort to incorporate into any existing enzymatic digestion protocols. Using a combination of our own snRNA-seq and a reanalysis of published literature we find that a similar signature is also expressed by post-mortem microglia and astrocytes. We also demonstrate that this post-mortem signature can be induced by technical variation in sample processing on human brain tissue, indicating that its presence in post-mortem brains may be artifactual. These datasets and methodologies should form the basis for future scRNA-seq/snRNA-seq experiments to avoid confounding impacts of technical signatures of identification and interpretation of true biological signals.

## Acknowledgments

Thanks to the Harvard Medical School’s O2 High Performance Compute Cluster.

## Author Contributions

S.E.M., A.J.W., L.D.O, T.R.H., and B.S. conceived the study. S.E.M., A.J.W., L.D.O, T.R.H., A.M.H.Y, S.H., E.M. and B.S. designed the experiments. S.E.M., A.J.W., L.D.O, T.R.H., C.D., and S.H. performed mouse cell isolation, sorting, and cell capture. A.M.H.Y., H.B., P.J.H., and R.J.M.F provided acutely-resected human tissue. A.A., N.N., and C.V. performed human nuclei isolation, sorting, and nuclei capture. S.E.M., A.A., and N.N. generated scRNA-seq/snRNA-seq libraries. S.E.M. performed library NGS sequencing. S.E.M., T.R.H., C.D., and S.M. performed smFISH experiments. S.E.M. and T.K. performed scRNA-seq/snRNA-seq analyses with assistance from A.J.W., T.R.H, V.K., and E.M. S.E.M. and B.S. wrote the paper with input from all authors.

## Methods

### Animals

C57BL/6J (Stock # 000664) were purchased from Jackson Laboratories. All mouse single cell sequencing studies were performed with male mice at P89-90. Both male and female mice were used for smFISH studies. All experiments were reviewed and overseen by the institutional animal use and care committee at Boston Children’s Hospital in accordance with all NIH guidelines for the humane treatment of animals. For full breakdown of replicates per dataset for mouse experiments see **SI Table 1**.

### Acute Human Tissue

Acutely isolated human brain tissue was obtained with informed consent under protocol REC 16/LO/2168 approved by the NHS Health Research Authority. Samples were transferred subject to MTA agreements between institutions and use and processing of acute brain tissue and sequencing were reviewed and approved by Boston Children’s Hospital Institutional Review Board and Broad Institute’s Office of Human Research Subjects Protection.

Adult brain tissue biopsies were taken from the site of neurosurgery resection for the original clinical indication. Samples were dissected into equal volumes and half was immediately (<5 minutes from time of tissue extraction) snap frozen in liquid nitrogen and stored at -80°C. For examination of technical component to gene expression profiles, the remaining half of the samples, were placed in Hibernate A low fluorescence (HALF) supplemented with 1x SOS (Cell Guidance Systems), 2% Glutamax (Life Technologies), 1% P/S (Sigma), 0.1% BSA (Sigma), insulin (4g/ml, Sigma), pyruvate (220 g/ml, Gibco) at room temperature for 2 hours and then 4°C for 4 hours before being snap frozen in liquid nitrogen and stored at-80°C. Samples were shipped on dry ice and stored at -80°C before processing for nuclei isolation, sorting, and sequencing.

### Post-Mortem Human Brain

Post-mortem autopsy tissue from control cases was obtained from the Massachusetts Alzheimer’s Disease Research Center (MADRC) at Massachusetts General Hospital (MGH). Tissue were collected with informed consent of patients or their relatives and approval of Massachusetts General Hospital Institutional Review Board.

Human patient demographic information for tissue in the current study is provided in **Table S2**. Post-mortem tissue processing and sequencing experiments were performed at the Broad Institute and approved by Broad Institute’s Office of Research Subject Protection.

### Inhibitor Cocktail to Prevent Activation

Inhibitor stocks were reconstituted and stored as follows. For Actinomycin D and Anisomycin stocks were kept for no longer than 1 month following reconstitution.

Actinomycin D (Sigma-Aldrich; Cat#: A1410) was reconstituted in DMSO at stock concentration of 5mg/ml and aliquoted and stored at -20°C protected from light. Triptolide (Sigma-Aldrich; Cat #: T3652) in DMSO at stock concentration of 10mM and aliquoted and stored at -20°C protected from light. Anisomycin (Sigma-Aldrich; Cat#: A9789) was reconstituted in DMSO at stock concentration of 10mg/ml and aliquoted and stored at +4°C protected from light.

Three different buffer solutions were used at different steps in protocol, Perfusion Buffer (PB), Dissection Buffer (DB), and Digestion Buffer (DgB). The inhibitor cocktail was added to 3 different steps of the protocol as follows:

#### Perfusion Buffer (PB+Inhib)

HBSS (w/o Ca^2+^, Mg^2+^, and Phenol Red; ThermoFisher Scientific Cat#: 14175-145), Actinomycin D final concentration 5 ug/ml (1:1000 from stock), Triptolide 10μM (1:1000 from stock). Add inhibitors immediately prior to beginning perfusion and keep on ice protected from light. Make fresh for each experiment.

#### Dissection Buffer (DB+Inhib)

HBSS (w/o Ca^2+^, Mg2+, and Phenol Red), Actinomycin D final concentration 5 μg/ml (1:1000 from stock), Triptolide 10μM (1:1000 from stock), Anisomycin 27.1 μg/ml (1:368.5 from stock). Add inhibitors immediately prior to beginning perfusion and keep on ice protected from light. Make fresh for each experiment.

#### Digestion Buffer (DgB+Inhib)

Digestion buffer/Enzyme Mix of choice, Actinomycin D final concentration 5 μg/ml (1:1000 from stock), Triptolide 10μM (1:1000 from stock), Anisomycin 27.1 μg/ml (1:368.5 from stock). Only add inhibitors to digestion mix immediately prior to the addition of tissue for digestion.

### Single Cell Isolation (All CNS Cell Types)

Mice were anesthetized and perfused intracardially with ice cold HBSS with or without inhibitor cocktail in perfusion buffer. Brains were quickly dissected and meninges removed as completely as possible and whole brains were placed in dissection buffer on ice with or without inhibitors and kept covered to protect from light. After all perfusions were completed brains were quickly placed into sagittal adult mouse brain slicer matrix (Zivic Instruments; Cat #: BSMAS005-2) and sliced into 6 even sections.

Immediately following brain slicing, slices were added to Miltenyi gentleMACS C Tubes with and without inhibitor cocktail added to the digestion mixture. Samples were placed in Miltenyi gentleMACS OctoDissociator and the 37C_ABDK_01 protocols was run according to manufacturer’s instructions. Once program finished the samples were briefly spun according to Miltenyi protocol before being filtered through 70μm filter. Samples were washed with HBSS and spun to pellet cells. For isolation of all CNS cells, cell pellets were resuspended and overlaid with appropriate volume of Miltenyi Debris Removal Solution according to Miltenyi protocol. Debris was removed from top layer and solution was diluted with HBSS and spun to pellet cells. Cells collected following density gradient centrifugation were counted manually via hemocytometer for loading into 10X Chromium instrument.

### Single Cell Isolation (Microglia/Myeloid Cells)

Enzymatic dissociation of microglia from adult brain was performed identically to all CNS cells described above until the density gradient centrifugation step. After filtering cell pellets were resuspended in 40% Percoll (GE Healthcare) in HBSS. Samples were spun for 1 hour at 500 x g at 4°C. Pelleted cells were washed with HBSS and centrifuged 5 min at 500 x g at 4°C and resuspended in FACS buffer (0.5% BSA, 1mM EDTA, 1X PBS; sterile filtered).

Mechanical dissociation was performed as previously described ^14^, with the addition to +/-inhibitor solutions during perfusion, dissection, and Dounce steps. Perfusion and dissection were performed identically to description above. Following dissection brains were minced with scalpel and then Dounce homogenized 15-20 times with loose pestle and then 15-20 times with tight pestle, all while simultaneously rotating the pestle. Cell suspension was then passed through pre-wet 70μm filter. Percoll density centrifugation and resuspension in FACS buffer was then performed as described above.

An additional dataset of animals who received a tail vein injection of PBS 18 hours prior to sac was processed identically to the mechanical dissociation dataset described above.

### Fluorescent Assisted Cell Sorting (FACS)

All steps were performed on ice or using pre-chilled refrigerated centrifuge set to 4°C with all buffers/solutions prechilled before addition to samples. Cell suspensions were incubated for 20 minutes on ice with anti-CD16/CD32 to block Fc receptors (1:50; BioLegend Cat #: 553141) and with a viability dye; eFlour780 (1:1000; Thermo Fisher Scientific Cat# 50-169-66) to identify live cells. The following antibodies were then added to cell suspensions at 1:200 concentrations: CD11b-PE (BioLegend Cat#: 101208; Clone: M1/70), CD45-APC/Cy7 (BioLegend Cat#: 103116; Clone: 30-F11), CX3CR1-APC (BioLegend Cat#: 149008; Clone: SA011F11s). Samples were incubated with staining antibodies for 20 min at 4°C and then spun down for 5 min at 300g, before being resuspended in 500μl of FACS buffer.

Sterile 96-well plates were precoated with 200μl FACS buffer and chilled for 1 hour during sample staining. All but 5μl were removed and plates kept chilled on ice until samples ready to sort. In order to keep samples chilled during as much of protocol as possible and prevent contamination each sample sorted into single well of individual plate. Gating strategy for myeloid cell sort was as follows **(SI Figure 1b**: Live (live/dead eFluor780), cells vs. debris (FSC-A vs. SSC-A), singlets (FSC-H vs. FSC-A), CD45+/CD11b+. We intentionally set a liberal gate of any CD11b+/ CD45+ cells so as to not bias our scRNA-seq due to changes in cell surface receptor expression **(SI Figure 1b-c)**. CX3CR1 was excluded as parameter used for cell sort due to differences in staining between isolation methods as result of cleavage of extracellular epitopes by enzymes **(SI Figure 1d-e)**. A total of 12,000 myeloid cells were sorted on special order BD FACSAria II using 70μm nozzle with purity mode, at total speed of ∼10,000 events per second. Each sample took ∼5-10 minutes total to sort 12,000 live, double positive, single cells. After sorting plates were sealed with plastic cover and placed back on ice until proceeding to 10X single cell capture.

The MFI intensity of the CX3CR1 fluorescence (APC) of the CD45+ and CD11b+ population of cells were analyzed using FlowJo-V10, with statistical analysis performed using Prism 8.

During the sorting of the tail vein PBS-injected dataset, a clog occurred during the sort of the final animal in that dataset. The cells sorted prior to the clog were discarded and the sample was placed back on ice. Following cleaning of the sorter the sample was sorted a second time into fresh well. Consequently, the sample was transferred between room temperature and ice twice for a total time at room temperature of approximately 20 mins.

### Single Cell Partitioning & Library Generation

All mouse single cell sequencing was performed used 10X Genomics 3’ Single Cell Version 2 kits. For the experiments sequencing all CNS cell types, the volume of cell suspension containing 10,000 cells was calculated from manual hemocytometer cell counts and added to appropriate volume of nuclease free H2O according to 10X Genomics 3’ Single Cell Version 2 user guide.

For experiments with sorted myeloid cells the entire volume of sorted cells (∼17-20ul) was removed from well and added to appropriate volume of nuclease free H_2_O.

Cell suspensions were loaded onto Chromium Single Cell Chip A with other RT reagents according to manufacturer’s protocol. Following droplet generation, barcoded single cell libraries were generated following manufacturer’s specifications. Library QC was performed using Agilent 2100 Bioanalyzer system using Agilent High Sensitivity DNA quantification kit (Cat #: 5067-4626).

### Single Nuclei Isolation & Sorting

Nuclei were isolated from human samples according to similar protocol previously published for use with mouse brain tissue ^73^ (https://www.protocols.io/view/frozen-tissue-nuclei-extraction-for-10xv3-snseq-bi62khge). See protocols.io link for all buffers and solution concentrations. All steps were performed on ice or cold block and all tubes, tips, and plates were precooled for >20 minutes prior to starting isolation. Briefly, 60 um sections of cortex (∼50 mg) were placed into a single well of six well plate and 5-6 ml of Extraction Buffer (ExB) was added to each well. Mechanical dissociation was performed through trituration using a P1000 pipette, pipetting 1ml of solution slowly up and down with a 1ml Rainin (tip #30389212) without creating froth/bubbles a total of 20 times. Tissue was let rest in buffer for 2 minutes and trituration was repeated. A total of 4-5 rounds of trituration and rest were performed (∼10 minuets). The entire volume of the well was then passed twice through 26-gauge needle into the same well. Following observation of complete tissue dissociation, ∼5-6 ml of tissue solution was transferred into a precooled 50 ml falcon tube. The falcon tube was filled with Wash Buffer (WB) to make the total volume 50mls. The 50ml of tissue solution was then split across 4 different 50 ml falcon tubes (∼12-13 ml of solution in each falcon tube). The tubes were then spun in a precooled swinging bucket centrifuge for 10 minutes, at 500 g at 4°C. Following spin, majority of supernatant was discarded (∼100μl remaining with pellet). Tissue solution from 4 falcon tubes were then pooled into single tube of ∼500μl of concentrated nuclear tissue solution. Approximately 500μl of WB was then added to bring the total volume of nuclei solution to 1 ml, in an Eppendorf tube. DAPI was then added to solution at manufacturer’s (ThermoFisher Scientific, #62248) recommended concentration (1:1000).

Flow sorting of isolated nuclei was performed similarly to protocols.io protocol (above). Briefly, 0.2ml PCR tube was coated with 5% BSA-DB solution. Solution was then removed and 20ul of FACS Capture Buffer was added as cushion for nuclei during sort. Nuclei were sorted into a chilled 96 well FACS plate (Sony M800 FACSorter). Sorting was done at a pressure of 6-7, with forward scatter gain of 1% on DAPI gate. The “purity” mode was used, and no spinning was performed after flow sorting nuclei into PCR tube. Following sort, nuclei concentration was counted using hemocytometer before loading into 10X Genomics 3’ V3 Chip.

### Single Nuclei Partitioning & Library Generation

All single nuclei experiments were performed using 10X Genomics 3’ Single Cell Version 3. Following droplet generation barcoded libraries were generated following manufacturer’s specifications with the following experiment specifics steps. To ensure sufficient numbers of microglia for downstream analysis we generated two libraries from each of the post-mortem samples.

### Next Generation Sequencing

All mouse single cell libraries were sequenced using Illumina NextSeq 500. All 16 mouse libraries (each representing individual animal for both all CNS cells and sorted myeloid cells) were diluted to according to Illumina specifications and combined into single library pool. Library pool was then prepared for sequencing according to Illumina denature and dilution guidelines for NextSeq500 High Output flow cells. Library pool was loaded on Illumina NextSeq 500/550 High Output v2.5 150 cycle flow cell and sequenced as follows according to 10X Genomics instructions for 3’ Single Cell Version 2 kits. Read 1: 26bp (16bp cell barcode, 10bp UMI); Index 1: 8bp (Illumina i7 sample index); Read 2: 98bp (Transcript insert).

The libraries from the tail vein PBS injected dataset submitted to Harvard Medical School’s BioPolymer’s sequencing core for quantification, pooling, and sequencing. Samples were QC’d via Agilent Tapestation and qPCR by Biopolymer Core before running on Illumina NextSeq 500/550 High Output v2.5 150 cycle flow cell and sequenced as follows according to 10X Genomics instructions for 3’ Single Cell Version 2 kits. Read 1: 26bp (16bp cell barcode, 10bp UMI); Index 1: 8bp (Illumina i7 sample index); Read 2: 98bp (Transcript insert).

Human single nuclei libraries were pooled to equimolar DNA concentration and sequenced at low depth (∼5000-10,000 reads/nucleus) on NextSeq500 in order to balance library pool by nuclei per sample before full depth sequencing using NovaSeq 6000. snRNA-seq libraries from both acute and post-mortem samples were pooled, diluted, and denatured according to Illumina specifications and loaded on Illumina NextSeq 500/550 High Output v2.5 150 cycle flowcell and sequenced as follows according to 10X Genomics specifications for 3’ Single Cell Version 3 kits. Read 1: 28bp (16bp cell barcode, 12bp UMI); Index 1: 8bp (Illumina i7 sample index); Read 2: 91bp (Transcript insert). Number of nuclei recovered per sample was determined via output of Cell Ranger ‘count’ pipeline (see below). Original libraries were then repooled to account for differences in nuclei number per sample to achieve equal read depth per nucleus across samples. New library pool was then sequenced on NovaSeq 6000 using S2 100 cycle flow cell using slightly modified read parameters (R1: 28bp, R2: 89bp, i7: 8bp). All NovaSeq 6000 sequencing was performed by the Broad Institute’s Genomics Platform.

### Single Cell/Nuclei Data Preprocessing

All preprocessing of sequencing data was performed on Harvard Medical School’s O2 High Performance Compute Cluster.

#### Mouse Data Preprocessing

Raw Illumina bcl files for the all CNS cells and sorted myeloid cells datasets were demultiplexed using Cell Ranger version 3.0.0 and bcl2fastq version 2.20.0.422 using ‘mkfastq’ step using default specifications. Individual sample gene expression matrices were generated using Cell Ranger version 3.0.0 ‘count’ step using default mm10 genome supplied by 10X Genomics Cell Ranger 3.0.0 (corresponds to filtered version of Ensembl v93; see 10X support website for further information). Sample specific results were then aggregated into combined output matrix using Cell Ranger version 3.0.0 ‘aggr’ function, specified with ‘normalize=none’ so that all reads for all samples were included in downstream analysis. Cell Ranger ‘aggr’ was performed individually for all CNS cell samples (n=4) and FACS sorted microglia samples (n=12).

The PBS tail vein injected dataset was processed using Cell Ranger version 2.2.0 and bcl2fastq version 2.20.0.422 using ‘mkfastq’ step using default specifications. Individual sample gene expression matrices were generated using Cell Ranger version 2.2.0 ‘count’ step using default mm10 genome supplied by 10X Genomics Cell Ranger v2.2.0 (corresponds to filtered version of Ensembl v84; see 10X support website for further information). Sample specific results were then aggregated into combined output matrix using Cell Ranger version 2.2.0 ‘aggr’ function, specified with ‘normalize=none’ so that all reads for all samples were included in downstream analysis.

#### Human Data Preprocessing

Raw Illumina bcl files were demultiplexed using Cell Ranger version 3.1.0 and bcl2fastq version2.20.0.422 using ‘mkfastq’ step using default specifications. Custom premRNA reference genome was generated using instructions from 10X Genomics. The default 10X GRCh38 genome (corresponds to filtered version of Ensembl v93) was used for modification. Individual sample gene expression matrices were generated using Cell Ranger version 3.1.0 ‘count’ step using this custom GRCh38 genome. Sample specific results were then aggregated into combined output matrix using Cell Ranger version 3.1.0 ‘aggr’ function, specified with ‘normalize=none’ so that all reads for all samples were included in downstream analysis. Cell Ranger ‘aggr’ was performed once for all human samples (n=10).

### Single Cell Analysis Tools

Mouse single cell analysis was performed using R (v3.4.3, v3.5.1, and 3.6.1) and primarily using the single cell analysis package Seurat (v2.3.4 and v3.1.5) ^74,75^. Additional analysis of human single nuclei samples was performed using LIGER development branch “online” ^37,38^. Other supplemental R packages were used as described in methods below.

Code required to reproduce Seurat or LIGER objects used for analyses and plotting can be found at: https://github.com/samuel-marsh/Marsh_et-al_2020_scRNAseq_Dissociation_Artifacts. Questions or correspondence regarding analysis/code can be directed to.

### Mouse scRNA-seq Analyses

Similar basic analysis pipelines were used in many of the analyses. Full details to replicate the analysis pipelines described briefly below can be found in code scripts available on GitHub.

### Initial QC & Clustering (Mouse Datasets)

In brief, cells were filtered using dataset specific parameters on basis of genes per cell, UMIs per cell, and percentage of mitochondrial gene reads per cell. Data was log normalized and highly variable genes were selected using mean expression and dispersion cutoffs. Data was scaled and UMIs per cell and mitochondrial gene percentage per cell were regressed out. Following PCA, relevant PC cutoffs were selected for downstream analyses based on combination of JackStraw analysis, ElbowPlot of PC variance, and manual examination of PCs. Clustering was then performed first by creation of KNN graph using PCs previously selected followed by Louvain clustering with dataset specific resolution parameter. Resolution parameter of Louvain clustering was iteratively performed to settle on appropriate final resolution. Dimensionality reduction was then performed using t-distributed Stochastic Neighbor Embedding (tSNE). Datasets were additionally QC’d at this step by checking for and removing doublets on basis of dual cell type marker gene expression. Following removal of doublets analysis pipeline was re-run using filtered dataset and slightly altered parameters.

### Cluster Annotation (Mouse)

Cluster annotation was performed through manual analysis of the output of differentially expressed marker genes from output of Wilcoxon Rank Sum test run via Seurat. Results were filtered in two different but complementary ways to identify marker genes, sorting for top genes by log fold change or by calculating a difference metric for percent of cells expressing vs. not expressing in given cluster and then sorting on top differences. For all CNS cell dataset marker genes were then used for annotation of cell type/subtype using previous single cell studies ^14,26,35,76,77^. For myeloid/microglia dataset clusters were annotated with names reflecting the genes enriched in that particular cluster (i.e. interferon-responsive, chemokine, etc) or on basis of known biology (i.e. proliferative).

### Differential Abundance Analysis (Mouse)

To determine if the proportions of cells per cluster were significantly different between the different protocols the clusters were compared using the R package speckle (https://github.com/Oshlack/speckle). The Seurat objects for the all CNS analysis and sorted microglia were used as inputs for the analysis. The design matrix for the all CNS analysis simply included the two experimental groups as comparison (Inhibitor vs. No Inhibitor). For the sorted microglia analysis in addition to the 4 experimental groups, the design matrix also included batch as variable.

### Subclustering (Mouse)

Subclustering analysis of all CNS cell types mouse dataset was performed by first calculating the number of cells per replicate for each of the clusters. Clusters were then combined into major cell classes by combining related/highly similar cell types. To enhance confidence in subclustering analysis only cell classes with greater than 80 cells per replicate (>300 cells total) were considered for subclustering. This criteria led to 6 cell classes for subclustering: 1) Myeloid (Microglia, exAM Microglia, Monocytes/Macrophages), 2) Astrocytes (Astrocyte 1, Astrocyte 2, Bergmann Glia), 3) Endothelial/Pericytes (Endothelial, Pericytes), 4) Oligodendrocytes (Oligodendrocytes and Oligodendrocyte Precursor Cells), 5) Epithelial (Epithelial/CP), 6) Neurons (Neurons and Neural Progenitor Cells). Each of the classes were subsetted from the original dataset and reanalyzed using similar pipeline to original analysis as described above.

### Metacell Differential Expression Analysis (Mouse)

To perform sample-level differential expression analyses we utilized the Metacell analysis as previously described ^14^. Each “meta-cell” is a pseudobulk replicate created by aggregating expression from all cells within a biological replicate. Pseudobulk profiles are then be analyzed via traditional RNAseq analysis pipelines. A recent comparative analysis found that pseudobulk analysis methods are among the best performing methods for differential state analysis and making sample-level comparisons from single cell data ^18^. For our pipeline, sample normalization and DE analysis was performed using DESeq2 ^19^ using simple pipeline. Genes were defined as differentially expressed with adjusted p value less than 0.05 and log2 fold change less than -0.58 or greater than 0.58 (corresponding to 1.5 fold change).

### Gene Module Scores (Mouse)

Creation of microglial identity scores and gene module scores were executed in Seurat based on previously published technique ^23^. Microglial identity score was based on well-established canonical microglial markers (**SI Table 3**). ‘Activation’ score was based on results of Meta-Cell pseudobulk differential expression analysis performed using DESeq2 **(SI Tables 3-10)**. For microglia/myeloid dataset consensus DE signature was identified by taking the intersection of the 3 pair-wise comparisons DNC-NONE vs. ENZ-NONE, DNC-INHIB vs. ENZ-NONE, ENZ-INHIB vs. ENZ-NONE.

### Re-analysis of publicly available datasets (Mouse Myeloid/Microglia)

Publicly available data, in the form of raw count matrices or loom files containing count matrices, were obtained from either from NCBI GEO, lab websites, or directly from authors as specified in **SI Figure 4g**. Information and links are also available via GitHub repository (see link above). For datasets made up of different genotypes or treatments only cells from WT or control mice were included in downstream analysis.

All datasets were processed with same basic Seurat pipeline with minor dataset specific changes (see code). Briefly, each dataset was filtered on dataset specific parameters for genes per cell, UMIs per cell, and percentage of mitochondrial genes. Data was log-normalized and variable features were selected using ‘mean.var.plot’ method using dataset specific thresholds. Data was then scaled and centered using all of the genes within the matrix for each particular dataset. Examination of PC loadings and results of JackStraw analysis were used to determine significant PCs for downstream clustering and visualization. Clustering was performed using the Louvain algorithm using dataset specific resolution parameters. Clustering results were visualized using tSNE. Activation and microglia scores were added as described above using the microglia Meta-Cell consensus DEG list.

For datasets containing more than just microglia/myeloid cells, the analysis pipeline was first run to identify myeloid cells. Clusters containing myeloid cells were isolated and entire pipeline run again to analyze myeloid cells using new parameters. For Zywitza et al., ^27^ analysis of isolated myeloid clusters revealed some astrocyte doublets which were removed following the second round of clustering and pipeline run a 3^rd^ and final time before analysis. For Mizrak et al., ^28^ analysis of isolated myeloid clusters revealed presence of astrocyte-microglia doublets, as well as some B and T cells. These cells were removed and the pipeline was run again for analysis of myeloid cells. For Zeisel et al., 2019 the microglia loom file was run through pipeline to generate new clustering. A small population of endothelial cells was found in the data during this initial clustering. These cells were removed the remaining cells were run through pipeline again.

### Human snRNA-seq Analysis

Similar basic analysis pipelines were used in many of the analyses. Full details to replicate the analysis pipelines described below can be found in code scripts available on GitHub (see link above)

### Initial QC Filtering (Human Datasets)

In brief, data was imported using Seurat V3 and QC filtered using dataset specific parameters on basis of genes per nucleus, UMIs per nucleus, and percentage of mitochondrial gene reads per nucleus. Datasets were then converted to LIGER objects with each sample serving as a separate “dataset” in LIGER.

### Doublet/Low Quality Nuclei Filtering (Human Datasets)

To detect and filter doublets and/or low-quality cells in snRNA-seq datasets we performed iterative rounds of clustering using LIGER. Most analyses utilized the new online iNMF algorithm ^38^ although some utilized traditional iNMF, when low cell numbers per sample were present. Following iNMF and quantile normalization, cells were clustered using Louvain clustering, followed by Uniform Manifold Approximation and Projection (UMAP) dimensionality reduction. Clusters were then annotated with one of the following broad cell class labels: excitatory neuron, inhibitory neuron, oligodendrocyte, oligodendrocyte progenitor cells (OPC), astrocyte, microglia, endothelial, fibroblast, pericyte, immune/PBMC (peripheral immune cells), or doublets. Annotation of broad cell classes and sub-types was performed using ^78^ and canonical marker genes as a reference. Each broad cell class was then subsetted and re-analyzed and clustered again. Doublet identification was then performed using combination of marker gene expression and shared LIGER ‘factors’ to identify nuclei that expressed markers or combinations of markers that are exclusive to two different cell classes. These subclusters were then classified as doublets and all barcodes corresponding to nuclei is those clusters were removed from the analysis. Some datasets required additional rounds of analysis and subclustering in order to completely remove likely doublet nuclei that were not found during the first round.

### Clustering and LIGER Factor Analysis

Following doublet removal, the “cleaned” subclusters for each major cell class were merged and full analysis and clustering was performed using LIGER. After cleaning, LIGER factors for each major cell class were examined by plotting the factor on UMAP plots and examining the top genes that loaded on each factor. Factors with genes indicative of activation, stress response, or similarity to mouse signature were selected for further analysis. For each selected factor we plotted the normalized cell-specific factor loadings for each gene and selected cutoff thresholds **(Figure 5a-b dashed lines)**. The list of genes above the cutoff in each factor (**SI Table 16)** was compared to the union of the two mouse DEG activation lists (**SI Table 3)** to analyze overlap and similarities.

### Gene Module Score Thresholding (Human Post-Mortem)

Quantify the enrichment of the microglia and astrocyte post-mortem LIGER factors across all samples in each dataset we performed gene module scoring using Seurat, as described above. To determine whether a particular cell’s score was defined as “enriched” we first determined dataset specific thresholds.

To determine thresholds for enrichment we first created an intersect gene list of all genes present across all 5 datasets. We then created 1000 random gene lists of equivalent length to the microglial LIGER factor (38 genes) and the astrocyte LIGER factor (27 genes). We then downsampled each dataset so that all of the major cell classes had the same number of cells, so that cell number/proportion did not influence outlier detection. We then performed gene module scoring using the random lists. To determine a cutoff for microglia we calculated an outlier threshold for each of the random scores. The outlier threshold was calculated as median score (across all cell types) + 3*median absolute deviation of score (across all cell types). We then plotted all of the outlier thresholds for each of the 1000 random scores. A cutoff was selected at the top 2.5% of outlier thresholds. We then determined the proportion of microglia per sample that exhibited a post-mortem LIGER microglia factor score above that cutoff. This process was repeated for astrocytes using the random gene lists of equivalent length to the astrocyte post-mortem LIGER factor. These enrichment proportions per sample were then used for correlational analysis to metadata variables (PMI, age, etc) **(Figure 5l-n, SI Figure 10a-c)**.

### Time Delay Freezing Analysis

Nuclei from acutely resected human tissue was analyzed using similar pipeline to previously described human tissue analysis. Following final clustering the microglial and astrocyte clusters were subsetted and converted to Seurat class for analysis comparing with post-mortem LIGER factors. Module scoring was performed using the LIGER factor genes **(SI Table 16)** as the input for postmortem microglial and astrocyte scores on the microglial and astrocyte subclusters respectively. DEG analysis was performed using Wilcoxon Rank Sum test comparing between the identity classes of 0hr vs. 6hr.

### RNAscope smFISH

Mice were anesthetized and perfused intracardially with ice cold HBSS. Brains were immediately dissected and flash frozen using vapor phase of liquid nitrogen. Following freezing brains were embedded in OCT (Tissue-Tek) and frozen on dry ice before storage at -80°C prior to sectioning. OCT embedded samples were mounted on cryostat and cut into 16 micron sagittal sections. Slides were kept frozen at all times during sectioning and then moved to -80°C for storage prior to RNAscope.

RNAscope Fluorescent Multiplex Assay (ACD Biosystems) was performed according to the manufacturer’s protocol for fresh-frozen tissue. Brains sections were hybridized with three mRNA probes per experiment. The following genes/ probes were used *Fcrls* (microglia/myeloid), *Ccl4*. The probes were amplified according to manufacturer’s instructions and labeled with the following fluorophores for each experiment: Alexa 488nm, Atto 550nm, Atto 647nm. High resolution images were taken using, 60x magnification using a Zeiss LSM XXX confocal microscope.

### Statistics

Statistical analysis of data was performed mainly using base R, Seurat, DESeq2, or speckle packages, with additional analyses in Prism 8 (GraphPad Software LLC).

### Data & Software Availability

Code required to reproduce Seurat or LIGER objects used for analyses and plotting can be found at: https://github.com/samuel-marsh/Marsh_et-al_2020_scRNAseq_Dissociation_Artifacts.

Raw sequencing data for all mouse samples was deposited in the NCBI GEO database under the SuperSeries GSE152184 which contains the following subseries: GSE152183 (Mouse Microglia 4 dissociation protocols), GSE152182 (Mouse All CNS Cells), GSE152210 (Mouse Microglia PBS Tail Vein). Cell Ranger output files are available as supplementary files via GEO and raw fastq files can be accessed from SRA linked from GEO records.

Raw sequencing data for post-mortem human tissue was deposited in NCBI GEO database under the super series GSE152184 in the subseries: GSE157760. Cell Ranger output files are available as supplementary files via GEO and raw fastq files can be accessed from SRA linked from GEO records.

Raw sequencing data for the acutely isolated human tissue was deposited in European phenome-Genome Archive (EGA) (Accession ID: *in progress*,).

