## Supplementary material for "Single Cell Sequencing Reveals Glial Specific Responses to Tissue Processing & Enzymatic Dissociation in Mice and Humans": SI Figures 1-11 & Legends

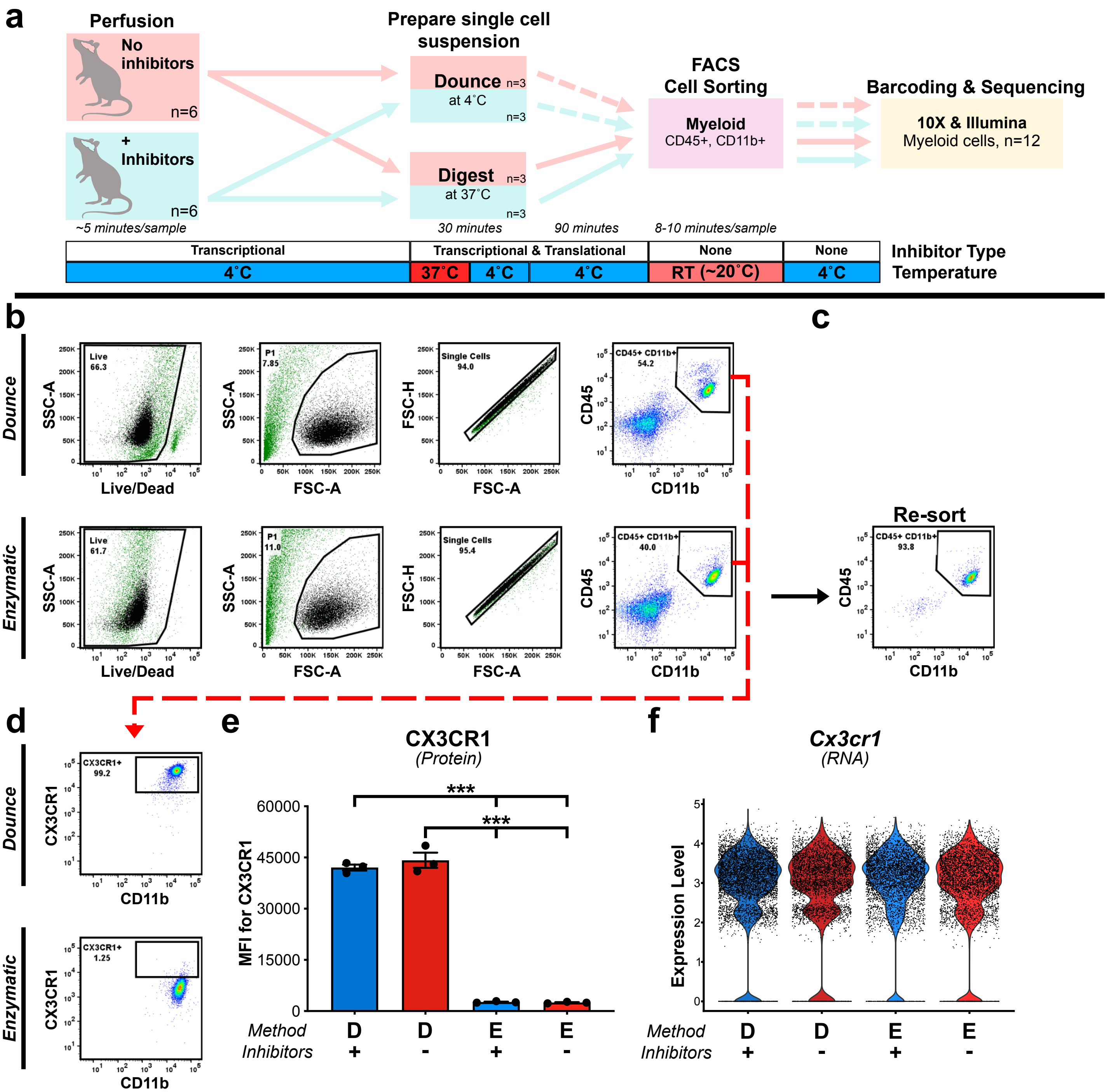


**Supplemental Figure 1: FACS gating strategy for single cell sequencing and differential protein expression based on digestion protocol. a.** Experimental design for sorted mouse microglia experiment (**Methods**). Mice were perfused with or without the inhibitor cocktail (n=6 per group). Groups were then further subdivided into enzymatic vs. mechanical Dounce digestion for total of 4 experimental groups (n=3/group). **b.** FACS gating strategy for scRNA-seq. Cells were first gated using Live/Dead stain, subsequently gated on overall size/granularity (FSC-A vs. SSC-A), and singlets (FSC-A vs. FSC-H). Cells were then sorted using dual positive CD45 and CD11b gate. **c.** Reanalysis of test sorted sample to examine purity of the sort. **d.** Flow plots from the sort gate showing expression of myeloid marker CX3CR1 vs. CD11b. **e.** Plot of median fluorescent intensity (MFI) for CX3CR1 across all samples. *** $p<0.0001$ of DNC-INHIB compared to either ENZ-INHIB or ENZ-NO-INHIB and DNC-NO-INHIB compared to either ENZ-INHIB or ENZ-NO-INHIB; two-way ANOVA with Tukey's multiple comparisons test post-hoc. **f.** Gene expression of *Cx3cr1* across all groups. **See SI Note 1** for discussion of CX3CR1 results and impact of enzymatic digestion on extracellular proteins.

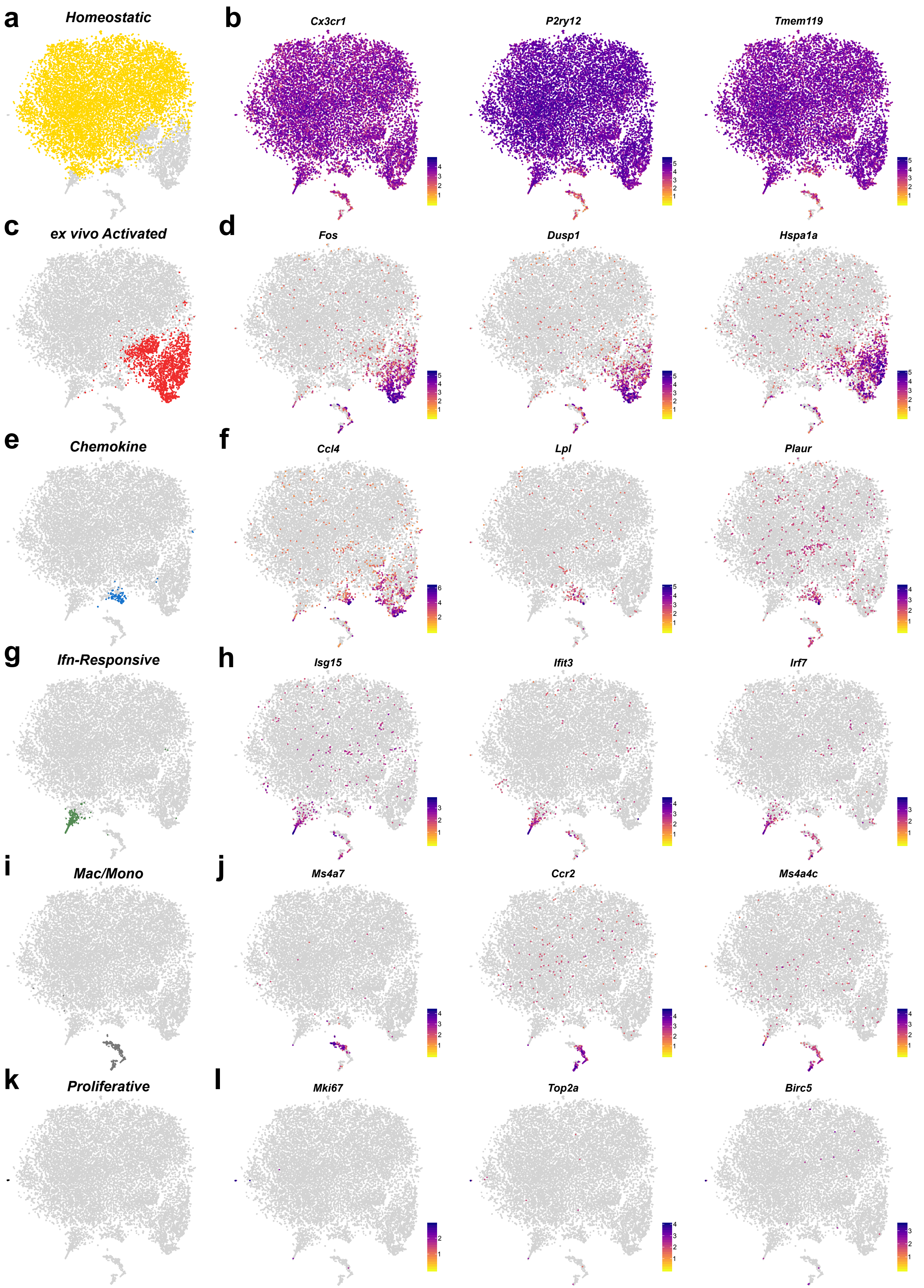


**Supplemental Figure 2: Cluster specific marker gene expression for mouse microglia clusters. a, c, e, g, i, k** tSNE plots with cells assigned to specific clusters, highlighted using same colors as in **Figure 1b**. **b, d, f, h, j, l** gene expression of cluster specific markers on tSNE coordinates.

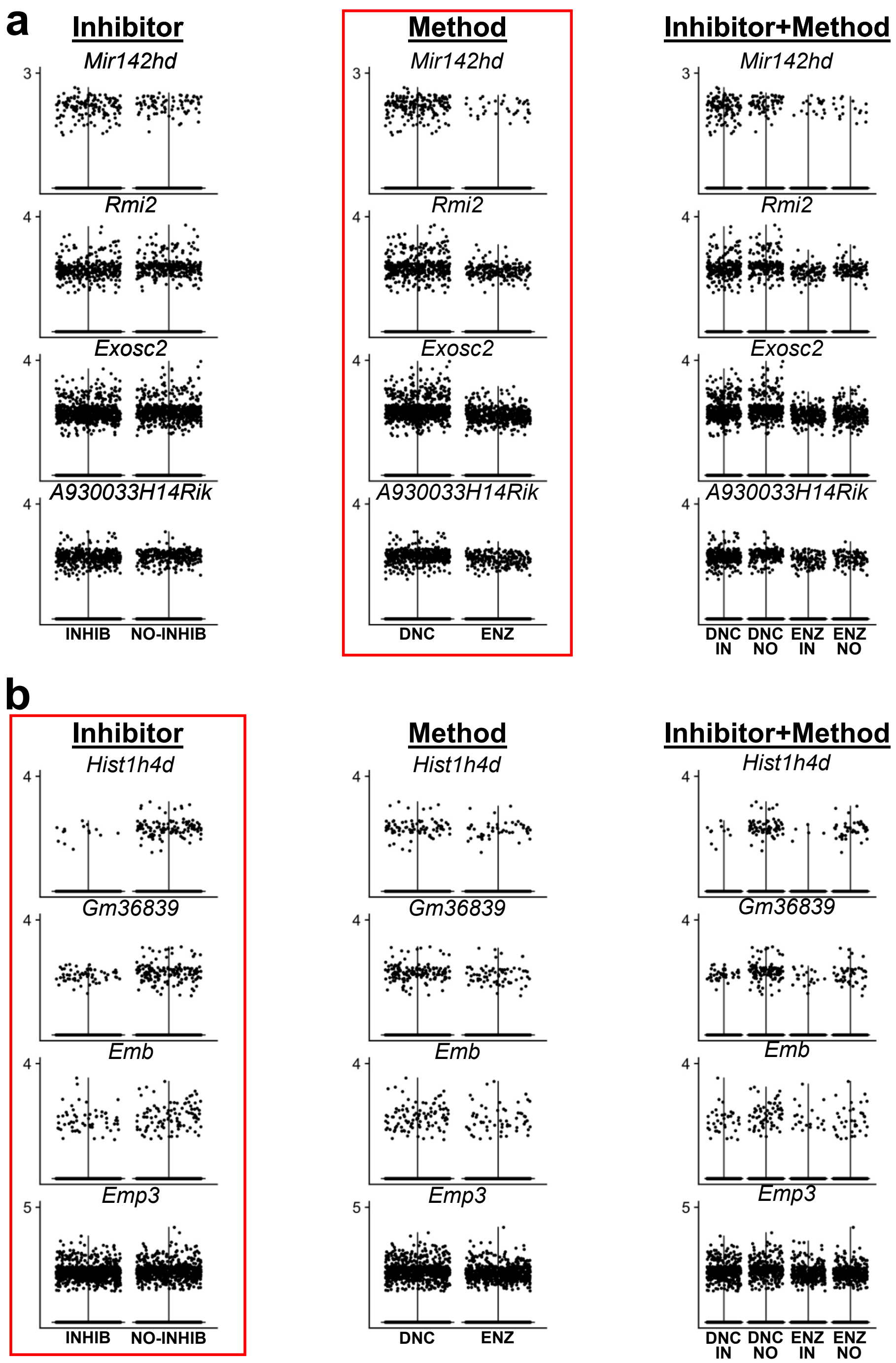


**Supplemental Figure 3: Analysis of potential negative consequences from use of inhibitors**. **a.** Gene expression values for a selection of genes from DE analysis comparison **(SI Table 5, 12)** showing primary effect is enzymatic digestion vs. Dounce dissociation. **b.** Gene expression values for a selection of genes from DE analysis comparison **(SI Table 12)** showing pattern with presence of inhibitors.

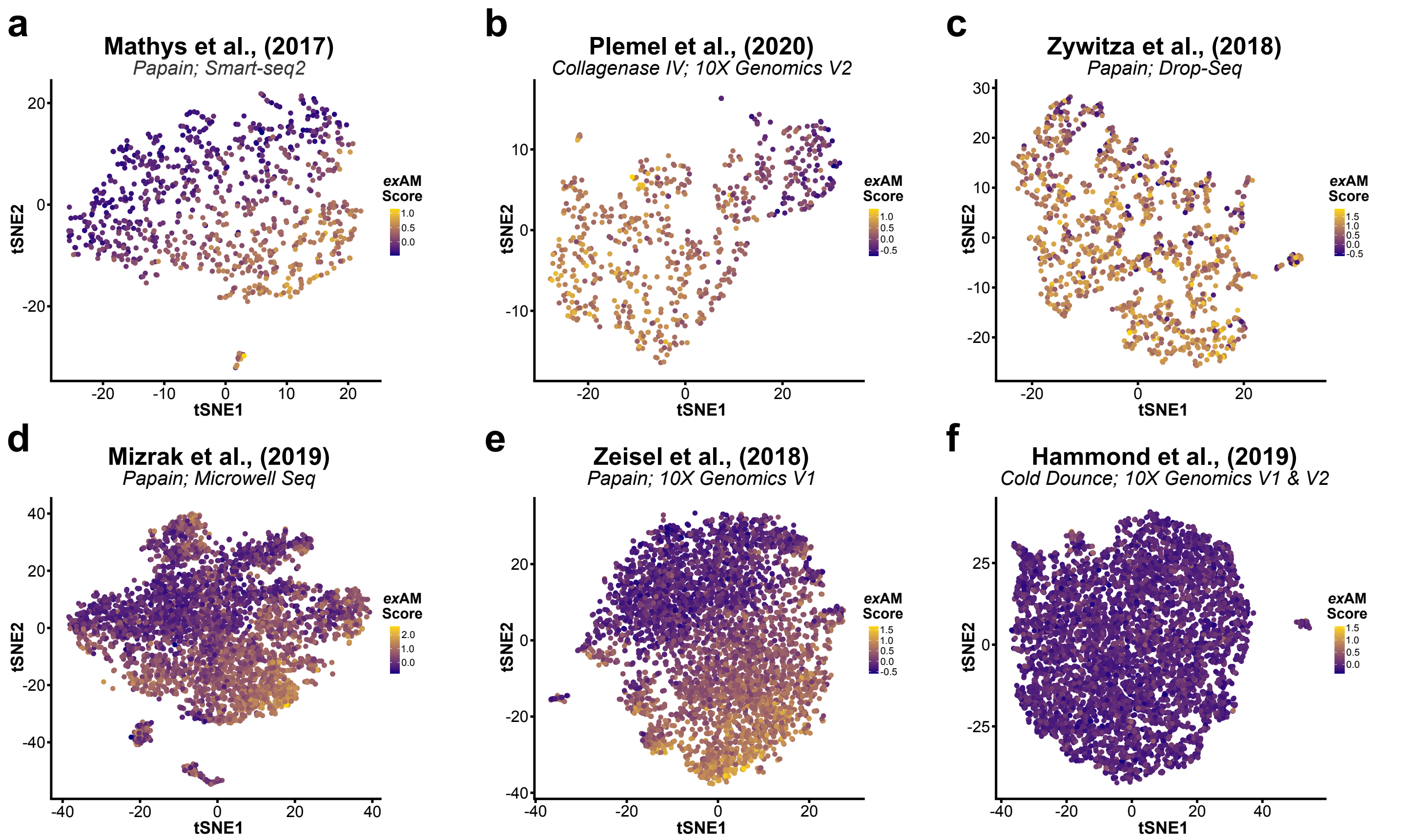


**g**

### Literature Reanalysis

Microglia Datasets

| Dataset Name | Enzyme | Seq Used | Tissue | Cell Number | GEO/SRA | Raw Counts Matrix | Citation |
|---|---|---|---|---|---|---|---|
| Mathys et al. | Papain (Miltenyi) | Smart-Seq2 | Brain | 883 | GSE103334 | Provided by Authors | Mathys et al. 2017 (Cell Reports) |
| Plemel et al. | Collagenase-IV | 10X 3' (V2) | Spinal Cord | 644 | GSE115803 | GEO | Plemel et al., 2020 (Science Advances) |
| Zywitza et al. | Papain (Miltenyi) | Drop-Seq | SVZ | 1153 | GSE111527 | GEO | Zywitza et al., 2018 (Cell Reports) |
| Mizrak et al. | Papain (Worthington) | Microwell Seq | SVZ | 5020 | GSE109447 | GEO | Mizrak et al., 2019 (Cell Reports) |
| Zeisel et al. | Papain (Worthington) | 10X 3' (V1 & V2) | Brain | 5227 | SRP135960 | mousebrain.org | Zeisel et al., 2018 (Cell) |
| Hammond et al | Dounce | 10X 3' (V1 & V2) | Brain | 7586 | GSE121654 | GEO | Hammond et al., 2019 (Immunity) |

**Supplemental Figure 4: *Ex vivo* artifactual gene expression occurs in brain myeloid populations when enzymatic digestion is used, across different labs, dissociation enzymes, sequencing technologies, and dissociation protocols.** **a-f.** Visualization of gene module scoring using consensus *ex*AM microglial activation gene module score **(SI Table 4-5)** projected onto tSNE coordinates. **a-e.** Reanalysis of brain myeloid cells from 5 sample datasets in the published literature, spanning variety of dissociation protocols, different dissociation enzymes, and five different sequencing technologies/technology versions. **f.** Reanalysis of our lab's previously published data with cold mechanical Dounce homogenization to present activation. **g**. Table of dataset characteristics and citations for the reanalysis in **a-f**.

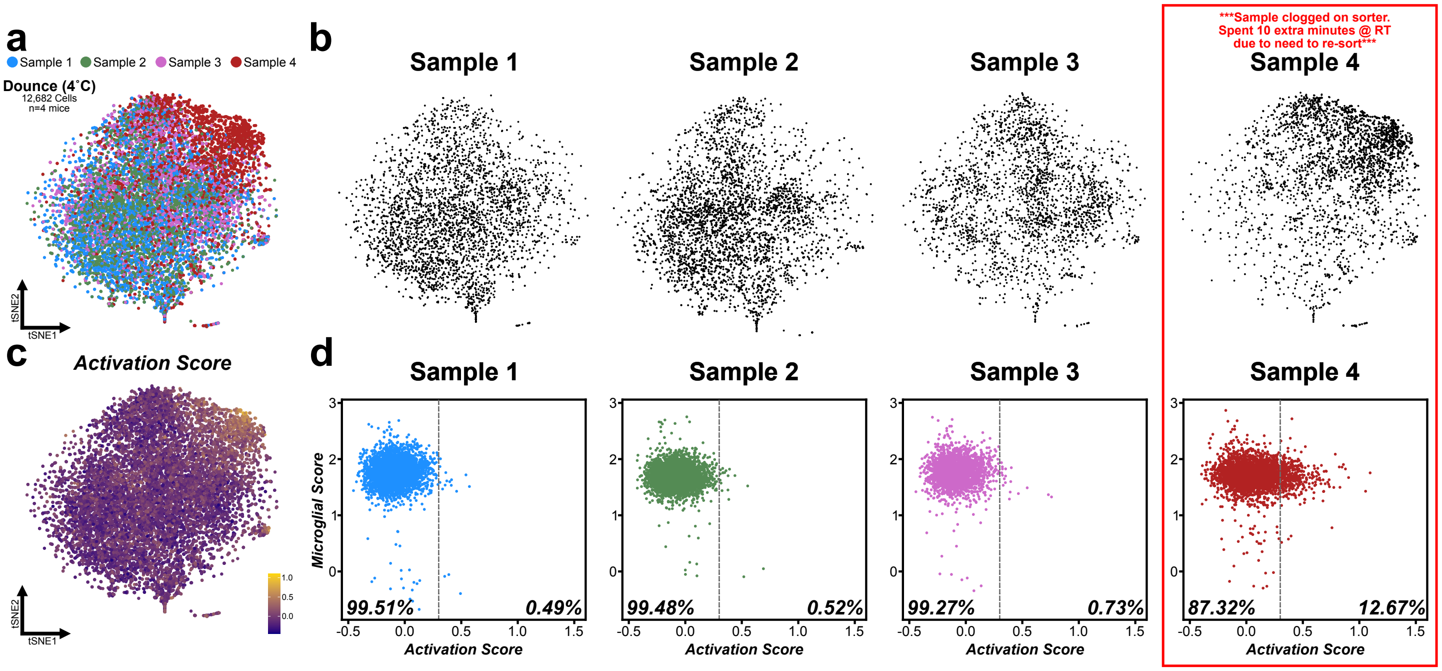


**Supplemental Figure 5: Other factors that can induce *ex vivo* alterations in gene expression even when using cold mechanical Dounce homogenization. a.** tSNE plot for an additional mouse microglia dataset of 12,682 cells from n = 4 mice that were injected with PBS via tail-vein 24 hours before sacrifice (control group), colored by replicate. **b.** Individual tSNE plot split by sample. **c.** Visualization of gene module scoring results using consensus *ex*AM microglial activation score **(see SI Table 4-5)** projected onto tSNE coordinates. **h.** Individual scatterplots by sample of the consensus *ex*AM activation score vs. the microglial identity score **(see SI Table 4)**. **See SI Note 2** for discussion of importance of biological replicates and controlling for variance even in optimized protocols.

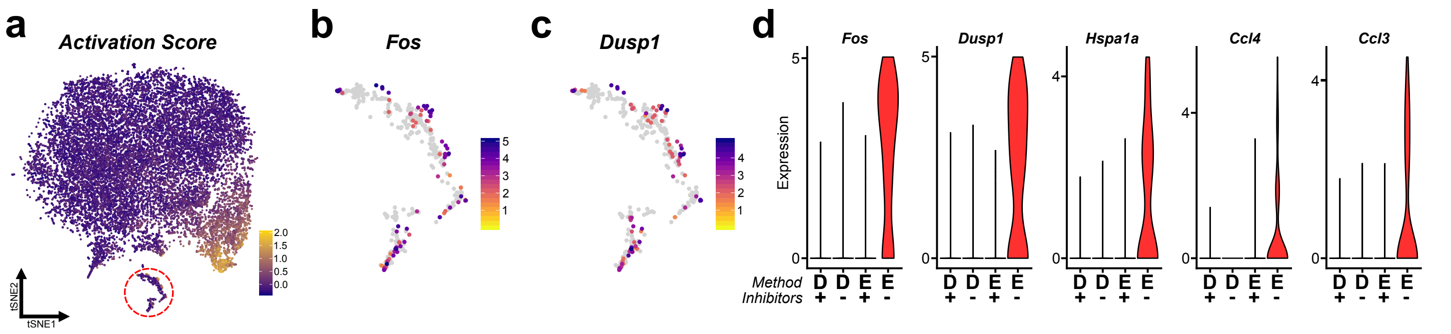


**Supplemental Figure 6: Non-microglial myeloid populations also exhibit *ex vivo* activation during enzymatic digestion. a.** tSNE plot of consensus *ex*AM signature in primary microglia/myeloid dataset (**Figure 1**), non-microglial myeloid cells highlighted (red dashed circle). **b-c.** Gene expression of *Fos* and *Dusp1* specifically in the macrophage/monocyte cluster circled in **a. d.** Gene expression of several *ex*AM markers split by experimental group demonstrate that increased expression of these markers only occurs in ENZ-NONE group.

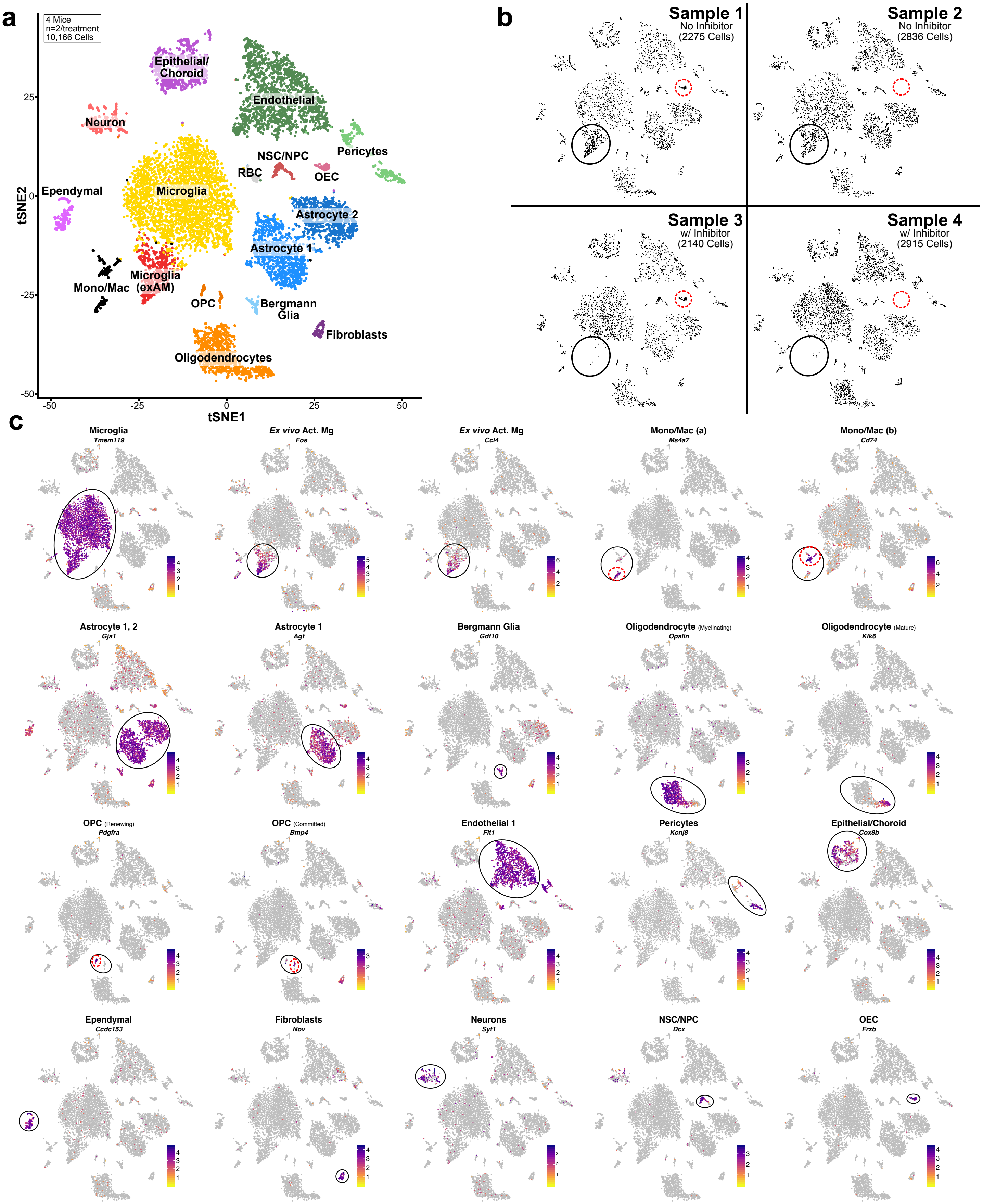


**Supplemental Figure 7: Identification of brain cell types from all CNS scRNAseq using enzymatic digestion with or without inhibitors. a.** tSNE plot of 10,166 cells from 4 mice, annotated and colored by cell type from (**Figure 2b**). **b.** tSNE plots for each of the 4 replicates (n = 2/group) individually. *ex*AM cluster location (black circle), OEC cluster location (red dashed circle). **c.** Cluster specific marker gene expression plotted on tSNE coordinates for all cell types and some subtypes.

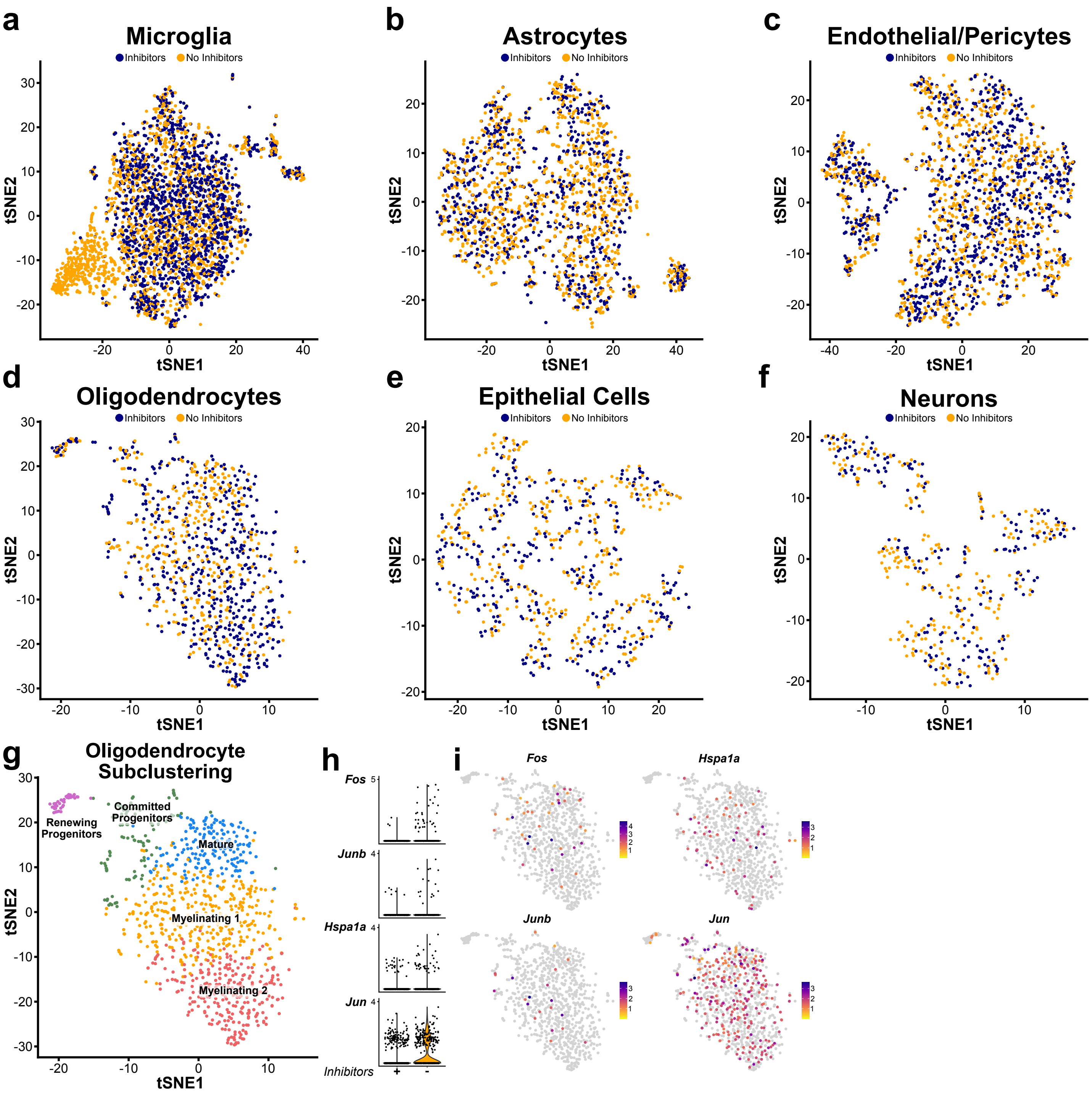


**Figure 8: Subclustering analysis confirms that microglia are particularly vulnerable to *ex vivo* artifactual gene expression during enzymatic dissociation. a-f.** tSNE plots from the subclustering analysis of major cell types present in **Fig 2b**, colored by presence/absence of inhibitors. **g.** Annotation of subclustered oligodendrocyte and oligodendrocyte progenitor populations. **h.** Gene expression for a selection of genes that are part of the activation score in oligodendrocyte subcluster split by experimental group. **i**. Gene expression of genes in **h** overlayed on tSNE coordinates for oligodendrocyte subclustering.

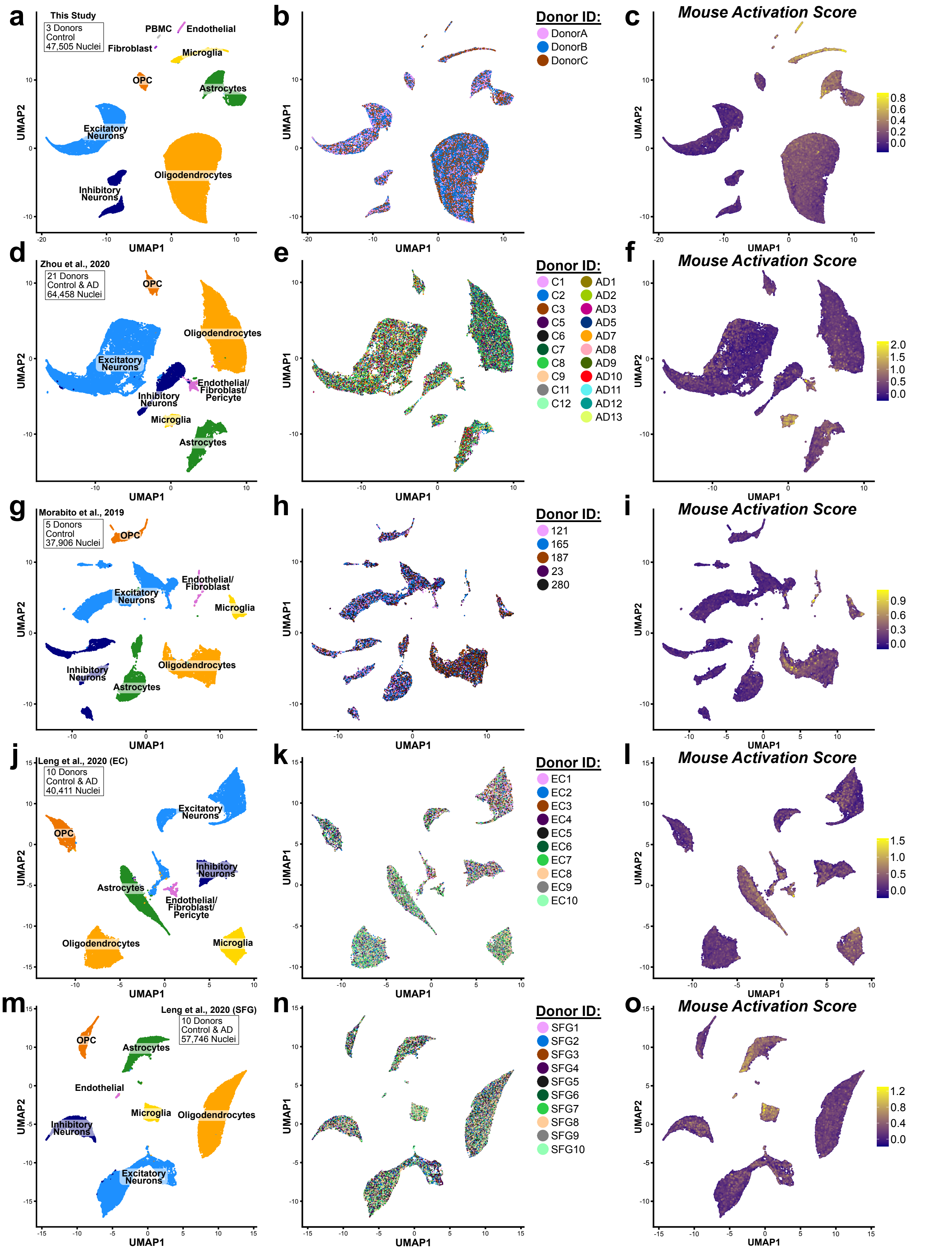


**Supplemental Figure 9: snRNAseq analysis and literature reanalysis of post-mortem human brain.** Total analysis includes 248,026 nuclei comprised of all major CNS cell types across 49 samples. For each dataset plots illustrate: overall cluster annotation by major cell type, integration of samples within dataset following LIGER analysis colored by donor, and visualization of gene module scoring using activation score from mouse dataset plotted on UMAP coordinates. **a-c.** Dataset produced in the current study; **d-f.** Dataset from Zhou et al., 2020; **g-i.** dataset from Morabito et al., 2020; **j-l.** dataset from Leng et al., 2020 (EC); **m-o.** dataset from Leng et al., 2020 (SFG).

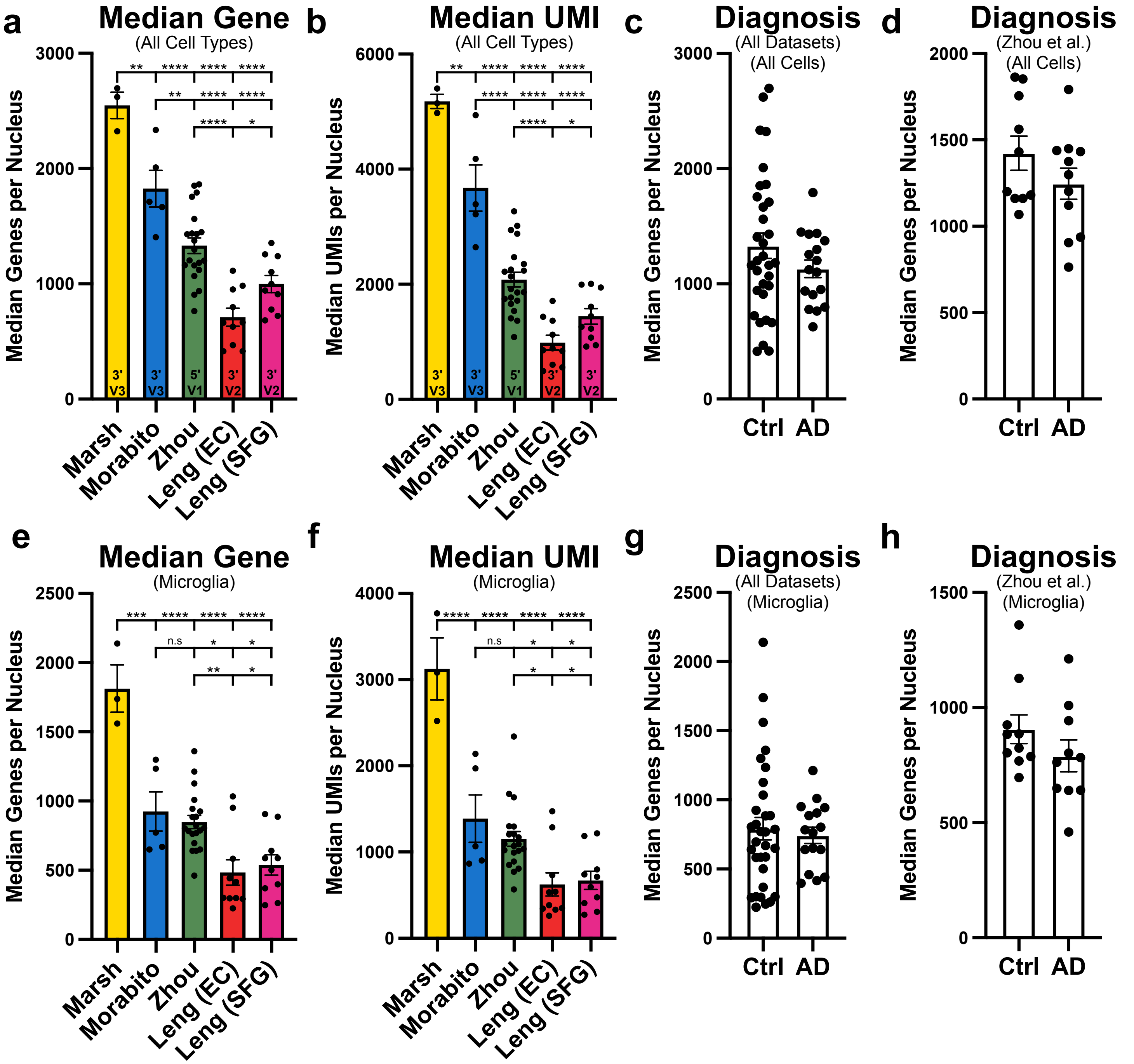


**Supplemental Figure 10: Significant differences in gene and UMI detection sensitivity across datasets demonstrates impact of 10X chemistry version and nuclei isolation protocol.** **a.** Median number of genes detected per nucleus for each sample across all cell types in each dataset. Version of 10X kit chemistry used in each study is listed inside bars of graph. **b.** Median number of UMIs detected per nucleus for each sample across all cell types in each dataset. Version of 10X kit chemistry used in each study is listed inside bars of graph. **c-d**. Median number of genes detected per nucleus for each sample across all cell types grouped by donor diagnosis for **c** all datasets or **d** just within Zhou et al., 2020 dataset. **e.** Median number of genes detected per nucleus for each sample in microglial nuclei in each dataset. **f.** Median number of UMIs detected per nucleus for each sample in microglial nuclei in each dataset. Version of 10X kit chemistry used in each study is listed inside bars of graph. **g-h**. Median number of genes detected per nucleus for each sample in microglial nuclei grouped by donor diagnosis for **g** all datasets or **h** just within Zhou et al., 2020 dataset. * $p<0.05$, ** $p<0.01$, *** $p<0.001$, **** $p<0.0001$; one-way ANOVA with Tukey's multiple comparisons test post-hoc.

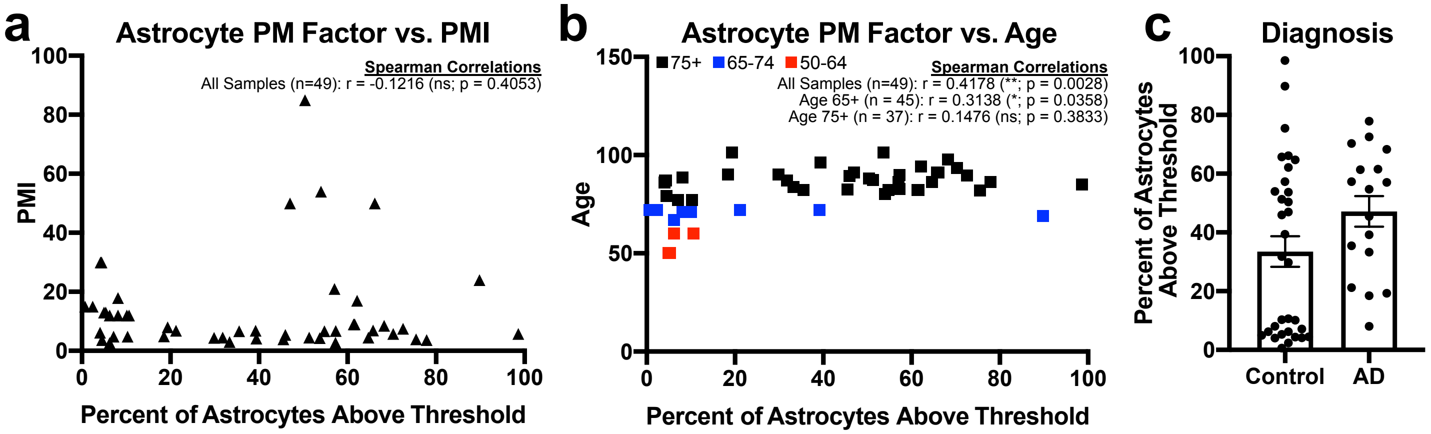


**SI Figure 11: Correlations between enrichment of astrocyte LIGER factor score and meta data variables in post-mortem astrocyte nuclei. a-b.** Correlation of PMI and age of donor vs percent of astrocytes above score threshold for astrocyte factor score in each post-mortem sample, graph annotations list Spearman r values and significance. **c.** Plot of percent of astrocytes above score threshold for astrocyte factor score in each post-mortem sample split by diagnosis.
