## Supplementary material for "Single Cell Sequencing Reveals Glial Specific Responses to Tissue Processing & Enzymatic Dissociation in Mice and Humans": SI Notes 1-3

### SI Note 1: Important considerations for the analysis of cell surface receptors/proteins following the use of enzymatic digestion (i.e. CITE-Seq, flow cytometry, CyTOF, etc)

In our previous study [1] we also used canonical myeloid marker CX3CR1 during the FACS sort. However, in the current study we noticed a dramatic reduction in CX3CR1 fluorescence in samples which had been enzymatically digested and therefore excluded CX3CR1 from the sort criteria (**SI Figure 1d-e**). Post-hoc analysis found that the presence of inhibitor cocktail during dissociation had no effect on CX3CR1, but cells digested with enzymes exhibited a nearly 17 fold decrease in median fluorescent intensity (MFI) of CX3CR1 (**SI Figure 1d,e**). A large body of work in the peripheral immunology field has demonstrated that enzymatic digestion is capable of cleaving extracellular epitopes from the cell surface [2–7], which is most likely what occurred in the current experiment as well. In addition to the short time scale of the experiment, RNA expression of *Cx3cr1* was stable across all four groups indicating no global downregulation of C*x3cr1* as a result of enzymatic dissociation (**SI Figure 1f**).

This potential caveat is further complicated by the fact that different enzymes have been found to result in cleavage of different epitopes with different affinities [2,3,5,6]. Understanding and controlling for the impact of enzymatic digestion is critical for any downstream method that measures extracellular protein levels. This is important for high-dimensional single cell-based methods that are used to complement scRNA-seq work (Flow cytometry, CyTOF, Infinity Flow) [8,9], but is perhaps most critical for high-dimensional multi-modal methods that pair examine of cell surface protein with transcriptomics (i.e. CITE-Seq) or epigenomics (i.e. ASAP-seq) [10,11]. Cleavage of extracellular epitopes by enzymatic digestion prior to CITE-Seq/ASAP-Seq may result in inaccurate linkage between RNA and protein levels for a particular gene/protein and analysis of such situations requires special care to control for these potential confounding effects. Similarly, results of comparisons made between cells from tissues digested with different enzymes [12,13] or between cells from undigested tissue (i.e. blood) vs. digested tissues [14] need to be interpreted with caution.

### SI Note 2: Importance of consistent tissue processing and biological replicates to enable identification of technical artifacts

While proper cold Dounce homogenization is sufficient to prevent *ex vivo* activation it is not infallible to experiment-specific. To demonstrate potential issues that may arise in course of normal experiment, we performed an analysis of a dataset we previously disregarded due to a technical issue that also used cold Dounce mechanical homogenization. This dataset consisted of the control group from an unrelated experiment (C57Bl6/J mice received tail vein injection of PBS 18 hours before perfusion and cell isolation). Following QC, this dataset consisted of 12,682 cells from n=4 mice. We noted that during the FACS sort of the final mouse (Sample 4) there was a clog in the sorter. Previously sorted cells were discarded and sample was placed back on ice while unclogging and cleaning was performed, before beginning the sort anew. This disruption meant that these cells went from ice to room temperature during first sort, back on ice, and then back to room temperature for the resort. Due to either extra time at room temperature, the extra set of temperature changes, or combination of factors we found that Sample 4 to exhibit significantly different characteristics compared to the first 3 samples.

When visualized by plotting the contribution of each sample to the overall tSNE plot, we observed an enrichment of Sample 4 in one corner of the plot (**SI Figure 5a-b**). To determine whether this enrichment represented *ex vivo* activated cells, we performed gene module scoring using the consensus *ex*AM signature. Plotting of this score revealed that the area enriched for Sample 4 was also enriched for cells positive for the activation signature (**SI Figure 5c**). We further confirmed that the cells exhibiting *ex*AM enrichment were almost exclusively limited to Sample 4 (**SI Figure 5d**). This finding demonstrated that even what might be considered minor deviations during cell isolation and sorting, can induce significant alterations in the transcriptome of microglia. This result also reinforces the need for well powered scRNA-seq experiments, including the need for independent biological replicates for each condition being examined, which is not always the case in the current scRNA-seq literature. Without proper replicates experiment-specific variations, such as the one described here, could be incorrectly attributed to the condition/treatment/genotype being examined as opposed to technical/batch effect

**SI Note 3: Substantial differences across published snRNA-seq literature in gene/transcript sensitivity likely corresponds to improvements/differences in single cell chemistry and nuclei isolation protocols.**

During our re-analysis of the literature, we did find that there were significant differences in dataset quality/sensitivity, across all cell types and on cell-type specific level **(SI Figure 10)**. The differences in genes/nucleus and UMIs/nucleus was significantly driven by both improvements in 10X Genomics chemistry across newer versions of their gene expression kits and differences between nuclei isolation protocols. Comparison of our results to that of Morabito et al., 2020 found that despite utilizing the same 10X 3' V3 chemistry and control neuropathology tissue with comparable RIN values (**SI Table 2**), we observed that on average our 39.5% greater median genes per nucleus and 40.9% greater UMIs per nucleus **(SI Figure 10a-b)**. When analyzing just microglial nuclei from each dataset the differences were even starker as on average our median gene per microglia nucleus was nearly double that of Morabito et al., 2020 (1813 vs. 925) and median UMI per microglia was nearly 2.5 times greater (3124 vs. 1389) **(SI Figure 10e-f)**. To determine whether donor diagnosis had any effect on gene or UMI detection we also compared across control and AD patients. When comparing either the entire dataset or just microglia we found no differences in between control and AD patients **(SI Figure 10c, g)**. To ensure that the lack of difference was not due to differences in 10X kit chemistry across datasets we also compared internally within the Zhou et al., 2020 dataset and found no differences as a result of diagnosis **(SI Figure 10d, h)**.
