## Supplementary material for "Single Cell Sequencing Reveals Glial Specific Responses to Tissue Processing & Enzymatic Dissociation in Mice and Humans": SI Tables 1-18: SI Table List_vInitial.docx

**SI Table 1:** Sample Information (Mouse)

**SI Table 2:** Sample Information (Human)

**SI Table 3:** Differential Abundance (Mouse Microglia)

**SI Table 4:** Gene Module Scoring Gene Lists

**SI Table 5:** Microglia Meta Cell Individual Comparisons Summary

**SI Table 6:** DESeq2 Sig Genes Results DNC-NONE vs. ENZ-NONE

**SI Table 7:** DESeq2 Sig Genes Results DNC-INHIB vs. ENZ-NONE

**SI Table 8:** DESeq2 Sig Genes Results ENZ-INHIB vs. ENZ-NONE

**SI Table 9:** DESeq2 Sig Genes Results DNC-NONE vs. DNC-INHIB

**SI Table 10:** DESeq2 Sig Genes Results DNC-NONE vs. ENZ-INHIB

**SI Table 11:** DESeq2 Sig Genes Results DNC-INHIB vs. ENZ-INHIB

**SI Table 12:** Effect of Inhibitors DESeq2 Overlap

**SI Table 13:** Overlap with other publications

**SI Table 14:** Differential Abundance (All CNS; Mouse)

**SI Table 15:** All CNS cells DESeq2 Sig Genes

**SI Table 16:** Post-Mortem Liger Factors Related to Mouse Signature

**SI Table 17:** Fresh 0hr vs. 6hr DEG Microglia Factor

**SI Table 18:** Fresh 0hr vs. 6hr DEG Astrocyte Factor
